# Identifying the environmental drivers of corridors and predicting connectivity between seasonal ranges in multiple populations of Alpine ibex (*Capra ibex*) as tools for conserving migration

**DOI:** 10.1101/2023.03.02.530594

**Authors:** Victor Chauveau, Mathieu Garel, Carole Toïgo, Pia Anderwald, Mathieu Beurier, Yoann Bunz, Michel Bouche, Francesca Cagnacci, Marie Canut, Jérôme Cavailhes, Ilka Champly, Flurin Filli, Alfred Frey-Roos, Gunther Gressmann, Ivar Herfindal, Florian Jurgeit, Laura Martinelli, Rodolphe Papet, Elodie Petit, Maurizio Ramanzin, Paola Semenzato, Eric Vannard, Anne Loison, Aurélie Coulon, Pascal Marchand

## Abstract

Seasonal migrations are central ecological processes connecting populations, species and ecosystems in time and space. Land migrations, such as those of ungulates, are particularly threatened by habitat transformations and fragmentation, climate change and other environmental changes caused by anthropogenic activities. Mountain ungulate migrations are neglected because they are relatively short, although traversing highly heterogeneous altitudinal gradients particularly exposed to anthropogenic threats. Detecting migration routes of these species and understanding their drivers is therefore of primary importance to predict connectivity and preserve ecosystem functions and services. The populations of Alpine ibex *Capra ibex*, an iconic species endemic to the Alps, have all been reintroduced from the last remnant source population. Because of their biology and conservation history, Alpine ibex populations are mostly disconnected. Hence, despite a general increase in abundance and overall distribution range, their conservation is strictly linked to the interplay between external threats and related behavioral responses, including space use and migration. By using 337 migratory tracks from 425 GPS-collared individuals from 15 Alpine ibex populations distributed across their entire range, we (i) identified the environmental drivers of movement corridors in both spring and autumn and (ii) compared the abilities of three modeling approaches to predict migratory movements between seasonal ranges of the 15 populations. Trade-offs between energy expenditure, food, and cover seemed to be the major driver of migration routes: steep south-facing snow-free slopes were selected while high elevation changes were avoided. This revealed the importance of favorable resources and an attempt to limit energy expenditures and perceived predation risk. Based on these findings, we provided efficient connectivity models to inform conservation of Alpine ibex and its habitats, and a framework for future research investigating connectivity in migratory species.

## INTRODUCTION

Global human-induced environmental changes are causing severe biodiversity loss, and habitat destruction and fragmentation are among the main causes of this decline (Newbold et al., 2016; Díaz et al., 2019). The development of linear infrastructures associated with human activities also contribute to impede species mobility (Torres et al., 2016). For instance, the extent of terrestrial mammalian movements was reduced by 50% in areas with a high human footprint compared with areas undisturbed by human activities (Tucker et al., 2018). By limiting animal movements between favorable habitats, human activities and infrastructures also reshape landscape connectivity (Taylor et al., 1993). Yet, connectivity is essential for individual and gene flows, for the local persistence of populations (Hanski, 1998), and for ecosystem functioning (Bauer & Hoye, 2014).

In the context of degraded connectivity, seasonal migrations, i.e., movements to track the spatiotemporal fluctuations in environmental conditions on seasonal ranges (Dingle & Drake, 2007), are of particular concern (Bolger et al., 2008). Most large herbivores, as primary consumers, migrate or may show migration propensity in heterogeneous and predictable habitats (Mueller at al., 2011, Teitelbaum et al., 2015). They are often restricted to well-defined corridors used by most migrants with low tendency to change migration routes when corridors are altered (see e.g., Xu et al., 2021). Migration can increase survival and reproduction through better access to high quality resources and reduced intra- and interspecific competition or predation risk (Avgar et al., 2014; Eggeman et al., 2016; van Moorter et al., 2021). However, migratory movements also imply energetic costs, and can be risky, or perceived as such, as animals may move through unfamiliar areas (Klaassen et al., 2014; Blagdon & Johnson, 2021). Hence, migration is a behavioral tactic whose fitness returns can vary through space and time, depending on individual traits, and spatial heterogeneity in occurrence and intensity of predation, harvesting, or competition in a population’s range. Accordingly, migration can be partial, with some individuals that choose to migrate while others are resident, and with individual behavior that can change from year to year (Cagnacci et al., 2011). Given that migration can affect population dynamics and species persistence by shaping their spatio- temporal distribution, there is a crucial need to increase our understanding of the link between habitat use and drivers of movement during seasonal migration at a fine spatial scale, the resulting ecological connectivity of a landscape, and how human activities affect this connectivity level (Sawyer et al., 2011; Panzacchi et al., 2016).

Migration corridors and their environmental characteristics are well-documented in spectacular collective and long-distance migrations in North American, Scandinavian or African ungulates (Boone et al., 2006; Merkle et al., 2016; Panzacchi et al., 2016; Joly et al., 2019) but are poorly known in most species (Kauffman et al., 2021). Recently, the focus has been put on spring migration revealing how migratory species can surf the green wave by tracking the green-up which moves like a wave across the landscape (Bischof et al., 2012; Merkle et al., 2016). Although less spectacular, migrations also occur in mountain ungulate populations occupying highly heterogeneous and fragmented landscapes (Herfindal et al., 2019), which are under threats from rapid climate changes and increasing anthropogenic pressure (Parmesan & Yohe, 2003; Schmeller et al., 2022). In mountain areas, green waves occur along altitudinal gradients and therefore green wave surfing seems to not always fully explain the choice of routes traveled between seasonal home ranges (Gaudry et al., 2015; Herfindal et al., 2019; but see Semenzato et al., 2021 for seasonal tracking of the altitudinal green and senescence wave). Several other factors can affect migration routes, particularly in complex topographic landscapes. Indeed, in addition to the diversity of migratory portfolios, migration is most often partial and takes place among multiple winter and summer ranges (Crampe et al., 2007; Lowrey et al., 2020; Denryter et al., 2021) and, up to now, little is known about migration patterns and migration routes for these mountain populations. Yet, this information is essential to improve the conservation of migratory species (e.g., through the establishment of protected areas, or to inform landscape planning; Mccollister & Manen, 2010) and preserve the ecological functions and ecosystem services migratory species support (Semmens et al., 2011). In this context, the importance of reliable connectivity maps for the identification of realistic corridors has been stressed (Sawyer et al., 2011; Zeller et al., 2012). A deeper understanding of the link between fine-grain habitat use and movements has been particularly invoked and up-to-date algorithms have been developed and used to model connectivity while accounting from iterative decisions of animals trading off exploration and optimal use of their environment (Panzacchi et al., 2016; Goicolea et al., 2021). However, population-specific movement analyses and connectivity predictions may be difficult to generalize over species and contexts when relying on samples from a single population not always representative of the species/habitat. Multi-populational analyses may be crucial to extend population-specific knowledge to species conservation, but such comparative analyses remain particularly scarce (Urbano, Cagnacci & Euromammals consortium, 2021).

Here, we investigated migration routes in several populations of Alpine ibex *Capra ibex* across the Alps in order to model and predict connectivity between summer and winter ranges. This mountain species of high patrimonial value went almost extinct during the XX^th^ century and recovered a large distribution thanks to intensive reintroduction programs (>55 000 individuals distributed across 178 populations; Brambilla et al., 2020). Even though most populations stem from individuals that were naive to the areas they were introduced into, seasonal migrations seem to occur in most populations. These migrations appear short in distance between a low altitude wintering area and a higher summer range. Yet, populations are still poorly connected, and drastic bottlenecks and founder effects have resulted in a very low level of genetic diversity (Biebach & Keller, 2009). Thus, effective conservation of this species and its habitats would highly benefit from better knowledge of the landscape characteristics used by ibex during migration, and from an assessment of the connectivity offered by available habitats. Owing to a unique GPS telemetry dataset from 425 ibex and 15 populations across the entire distribution range of the species, we first aimed at determining the environmental drivers of migratory tracks accounting for the many factors likely at play in an individual’s choice of routes. We specifically tested whether individuals (i) minimized energy expenditures and difficulties to travel by avoiding elevation changes, rugged terrain and snowy areas as traveling costs are paramount in all optimality models aiming at understanding the costs and benefits of migration tactics (Holt & Fryxell, 2011), (ii) selected habitats offering food resources and refuge from perceived predation risk, (iii) used visual landmarks (linear features such as ridges, tree lines and valley bottoms) as “compasses’’ (Alerstam & Bäckman, 2018), and (iv) avoided proximity to anthropogenic infrastructures (roads and ski areas; Table 1) during migration. Then, we took advantage of the 15 populations monitored to compare connectivity assessments when using either population-specific habitat selection criteria or criteria averaged over all populations; and to perform external validation of the ability of our model to accurately predict ibex migratory connectivity in the absence of any data on ibex locations, a crucial step to provide reliable information for species migration conservation across its native range.

**Table 1:** Hypotheses tested in the integrated Step Selection Analyses and their corresponding predictions.

| *Hypotheses* | *Covariables* | *Predictions* | *References* |
| --- | --- | --- | --- |
| Ibex minimize energy expenditures and traveling difficulties | Total elevation change | Ibex should perform steps with relatively low total elevation change | Passoni et al., 2021 |
|  | Ruggedness | Ibex should avoid rugged terrain which tends to increase movement costs and reduce visibility | Wall et al., 2006; Halsey & White, 2017 |
|  | Snow cover index | Ibex should avoid snowy areas which impede ibex movements | Richard et al., 2014; Sheppard et al., 2021 |
| Ibex select areas that can provide forage, security and thermal shelters | Northness | Ibex should prefer south exposed terrain as they present snow-free areas with access to early growing vegetation in spring or thermal shelters in autumn |  |
|  | Proximity to refuges (steep slopes) | Ibex should stay close to steep slopes to reduce perceived predation risk | Grignolio et al., 2007; Iribarren & Kotler, 2012 |
|  | Forest | Ibex should avoid forests as they prefer open habitats, being primarily grass roughage eaters | Parrini et al., 2009 |
| Ibex use landmarks for orientation | Proximity to ridges | Ibex should select for proximity to ridges and follow ridges during their migration |  |
|  | Proximity to valley bottoms | Ibex should avoid to go down to valley bottoms in spring when migrating up the mountain but opposite in autumn |  |
|  | Proximity to tree lines | Ibex should use tree lines as a landmark to follow in some populations but not in others |  |
| Ibex avoid human-linear infrastructures | Proximity to roads | Ibex should avoid roads constituting physical barriers or because associated with humans | Seigle-Ferrand et al., 2022 |
|  | Proximity to ski areas | Ibex should avoid ski areas because associated with humans | Dickie et al., 2020 |

## MATERIALS AND METHODS

### Study areas and GPS data

We relied on a GPS dataset collected on 425 individual Alpine ibex (*Capra ibex*; 41% females and 59% males; 77% being adults >4 years-old) from 15 reintroduced populations. Those populations were distributed across the whole Alps (10 in France, 2 in Italy, 1 in Switzerland, and 2 in Austria; Figure 1; latitudinal gradient: 44-47°N, longitudinal gradient: 6°-13°E and altitudinal gradient: 1700-2700m). Alpine ibex can share habitats with northern chamois (*Rupicapra rupicapra*), less frequently with red deer (*Cervus elaphus*) and roe deer (*Capreolus capreolus*), and with livestock during summer (sheep, goats and cows). Grey wolf (*Canis lupus*) is present in most of ibex distribution range, but predation occurs rarely on ibex.

**Figure 1:**
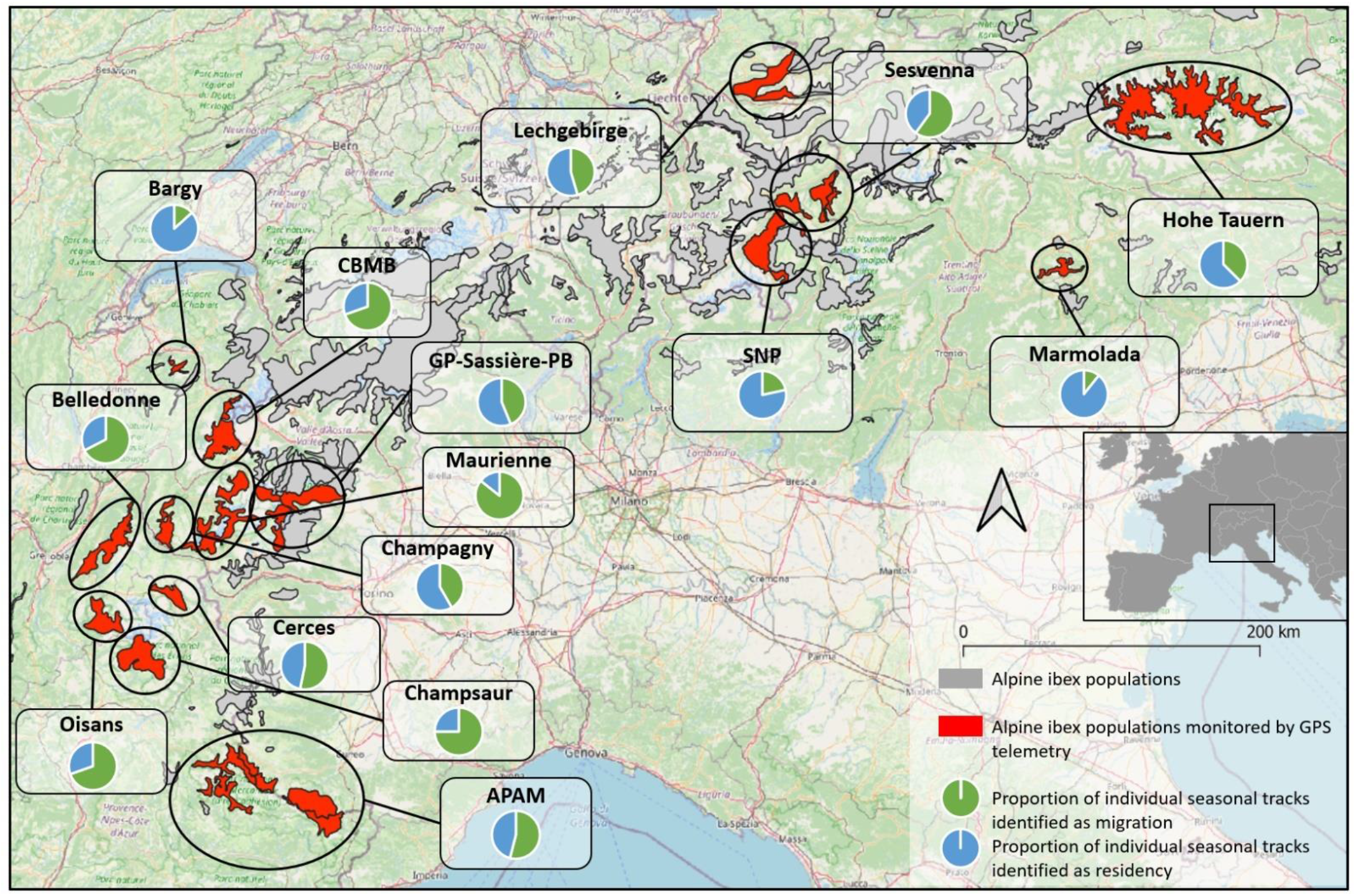
Location of the 15 Alpine ibex *Capra ibex* populations monitored by GPS telemetry (red) over the distributional range of the species (gray; source: Brambilla et al., 2020); see Appendix S1 for more details. The pie charts display the proportion of individual seasonal tracks identified as migration (green) or residency (blue) within each population (see below and Appendix S3 for details on individual status identification).

Sample sizes varied between populations (minimum: 7 individuals in Hohe Tauern National Park; maximum: 117 in the Bargy massif). Several types of collars were used (Vectronic: GPS Plus, Vertex Plus, or Vertex Lite models; Lotek: 3300S or Litetrack models; Followit: Tellus model). All models weighted <3% of individual body weight. They were programmed to record ibex locations at variable frequencies and during variable periods (from 1 location per hour during one year to 1 location per 6 hours during 2-3 years, Appendix S1), resulting in 1 068 seasonal tracks (an individual monitored during 1 year resulted in 2 potential migratory tracks).

### Determining migratory status and migration tracks

The migratory status of each ibex (migrant or resident) and migration tracks (for migrants) were visually determined using the application Migration Mapper^TM^ (version 2.3, Merkle et al., 2022; see Appendix S2 for the parameters used). This application provides tools to visually identify migrants, migration periods and tracks using the Net Squared Displacement (NSD; squared Euclidean distance between the first locations of the GPS trajectory and the following one; Börger & Fryxell, 2012; Appendix S3). Spring and fall migration periods and migratory tracks started at the last location preceding the increase/decrease in the NSD and ended at the first location when the NSD stabilized. Migratory movements were identified irrespective of distance separating seasonal ranges as ibex exhibited several forms of migratory movements within populations (short-altitudinal movements or long-distance movements). However, only neat migrations (i.e., two movements between distinct seasonal ranges) were selected to reduce uncertainty in the displayed behavior (see Appendix S4 for details on distance and altitudinal interval between seasonal ranges of migrant and resident individuals).

### Assessing environmental drivers of ibex migratory movements

#### Environmental variables

We investigated the influence of 11 environmental variables (see Appendix S5) that could affect movement choices during migration (Table 1). We considered the total elevation change (sum of changes in elevation values along a step), the ruggedness (Vector Ruggedness Measure; Sappington et al., 2007) and a snow cover index (calculated as the total annual number of days a pixel was covered by snow) as metrics reflecting the energetic costs and difficulties to travel during migration. We used the northness (cosine of aspect derived from the same DEM) and the snow cover index to reflect the accessibility and the quality of vegetation resources and the presence of snow cover, as well as the availability of thermal shelters. Contrary to what is commonly done in studies on migratory ungulates, we did not use vegetation variables or derived metrics (NDVI, Instantaneous Rate of Green-Up; e.g. Bischof et al., 2012) as we judged the information given by the northness and snow cover more relevant considering the short duration and distance of ibex migrations (see results). As ibex mostly use open areas (Parrini et al., 2009) and often steep slopes as refuge from perceived predation risk and human disturbance, we expected forests to be avoided and proximity to slopes >40° to be selected during migration (Grignolio et al., 2007; Iribarren & Kotler, 2012). We considered ridges, valleys and tree lines as potential visual landmarks used for navigation (Alerstam & Bäckman, 2018). Finally, we hypothesized that the proximity to roads and ski areas (i.e., human infrastructures that occasionally occurred in the surrounding of ibex population ranges) would be avoided as both can constitute barriers - physical barriers such as roads - or perceived as such because associated with humans for both roads and ski areas.

#### Habitat selection analyses during migration

We used integrated Step Selection Analyses (iSSA; Avgar et al., 2016) to assess the environmental drivers of ibex migratory movements. The habitat variables along (total elevation change) or at the end (all other habitat variables) of each used movement step (considered as the straight lines between recorded ibex locations) traveled by one individual ibex during migration were compared with the habitat characteristics along/at the end of 15 available steps it could have traveled, using conditional logistic regressions (Fortin et al., 2005; Thurfjell et al., 2014). We generated those available steps by sampling step lengths (corrected to get three dimensional lengths using a DEM at a resolution of 25m, which are particularly relevant in mountainous landscapes) and turning angles in parametric distributions (gamma distribution for the step length and Von Mises distribution for the turning angles; Duchesne et al., 2015) derived from the observed step length and turning angle distributions of the used steps. We accounted for the variable step duration in our dataset by deriving specific distributions for each step duration and checked if habitat selection was similar for the different timesteps by using the method of Used Calibration Plots (see Appendix S6).

We scaled habitat variables across all populations (i.e., variables were centered and divided by their standard deviation) to make their effect size comparable in iSSA outputs and we checked for potential correlations between our variables using a Pearson correlation. Correlation coefficients were > 0.3 for the variables forest and proximity to forest (ρ = 0.5) and for the variables step length, log(step length) and total elevation change (ρ = 0.6 step length/log(step length), ρ = 0.6 total elevation change/log(step length), ρ = 0.9 step length/total elevation change).

The logistic regressions included the 11 environmental variables, step length, log of step length and cosine of turning angles (Avgar et al., 2016). We chose to include the step length and turning angles in iSSA without interactions with habitat variables to simplify our models, except for the interaction between step length and elevation change. We fitted one model for each season (i.e., spring or autumn migration) and each of the 15 populations. For 6 populations in spring and 5 in autumn, the variables “forest” and “proximity to ski areas” were excluded from models as forest or ski areas were rare or absent in the distributional range of those populations. We chose to fit models at population scale because we were more interested in modeling migratory movements at the population scale. Accounting for sex-specific differences can be important for a species like Alpine ibex knowing to exhibit different patterns of movements between sexes (Herfindal et al., 2019). However, numbers of migrant females (or even migrant animals) were too small in several populations to test sex-specific differences (see Appendix S7). We investigated if habitat selection results differed between sexes in Appendix S8. We fitted models using the *clogit* function from the package *survival* in R V. 4.2.2 (Therneau, 2022, R Core Team, 2022). We conducted a model selection based on AICc with the *dredge* function in package *MuMIn* (Bartoń, 2022). The coefficients from the best models, i.e., models with an ΔAICc <2, were averaged using the *model.avg* function in the package *MuMIn*. We did not include “individual” as a random effect as the models did not converge correctly when doing so. We finally produced Used Habitat Calibration plots (UHC plots; Fieberg et al., 2018) using mixed-effect Poisson models (Appendix S9) to check for the agreement between model predictions and values of our covariates observed at or along used steps (Appendix S6).

### Building and validating models of migratory connectivity in ibex

We proceeded in four steps (see below and step III of Figure 2) to build and validate connectivity models based on the 15 populations to perform three different validation procedures (i.e., using three different pairs of training/validation datasets; Figure 2) designed to understand how our models could inform different management measures. The first procedure (“leave 10% of whole data out”) seeks to understand if based on all information we have on ibex habitat selection we can predict migratory movements of non-marked individuals in monitored populations. We used the second procedure (“leave 10% of population data out) to assess if based on data from a limited number of animals in a population we can predict where other animals should migrate. Finally, the third procedure (“leave one population out”) evaluates if based on all information we have we can predict migratory movements in populations without monitoring.

**Figure 2:**
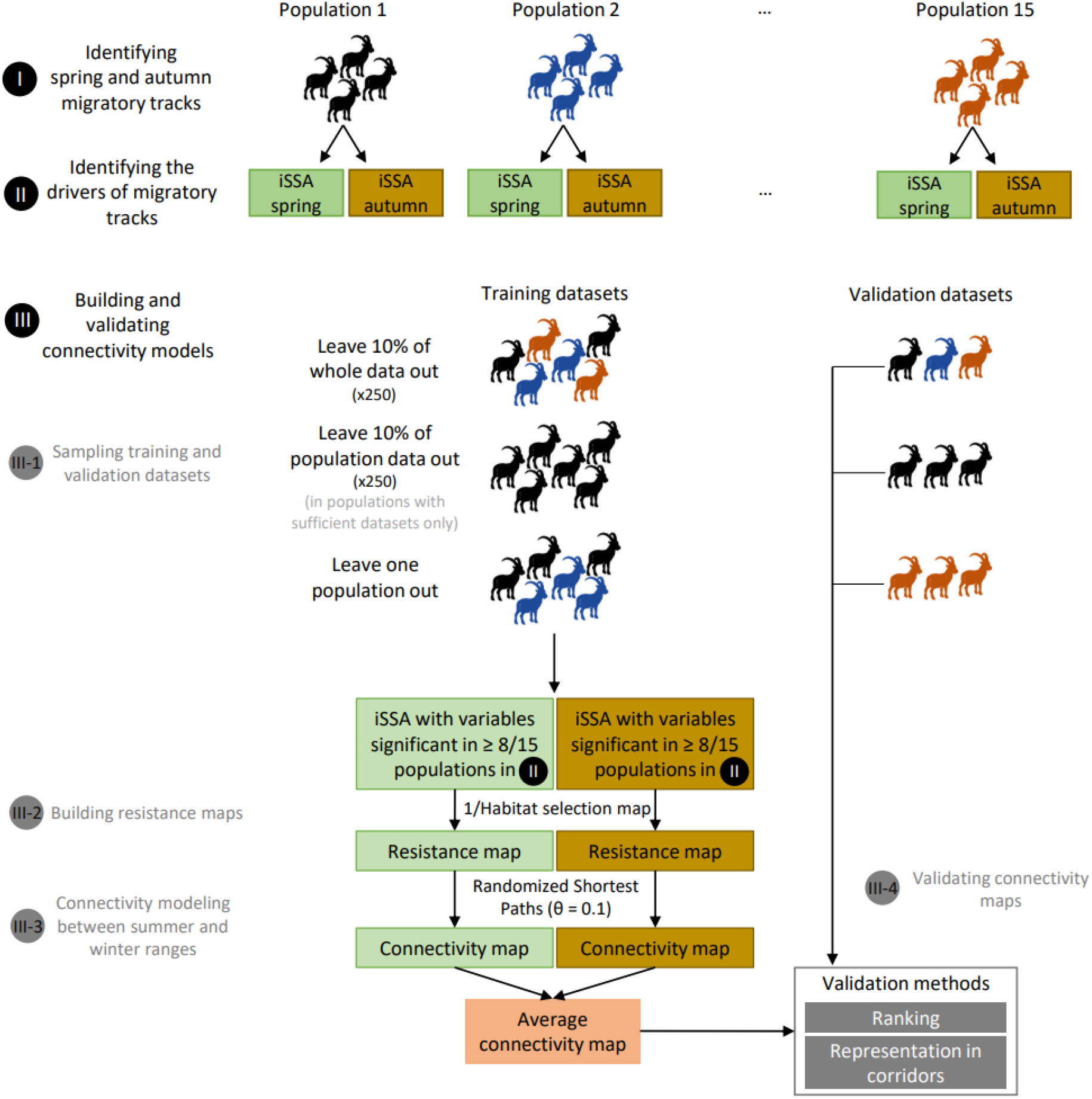
Methodological workflow scheme. First, we identified seasonal migratory tracks. Second, we fitted integrated Step Selection Analyses (iSSA) for each population to identify environmental drivers of spring and autumn migrations. Third, we sampled three different training and validation datasets: leave 10% of whole data out, leave 10% of population data out and leave one population out. The three training datasets were used in iSSA with the environmental variables identified as significant in at least 8/15 populations. We built three (one per training dataset) different resistance maps (250 times for “leave 10% of whole data out” and “leave 10% of population data out”) for each population and season and modeled connectivity based on these resistance maps using the Randomized Shortest Path algorithm (θ = 0.1; Appendix S10). Finally, for each training dataset, we averaged the two seasonal connectivity maps and combined this average connectivity map with the corresponding validation datasets to perform the ‘ranking’ ’and ‘representation in corridors’ validation methods.

For each seasonal migration (i.e., spring or fall), we first selected habitat variables previously identified as having a significant effect in at least 8/15 populations. To assess if variables had a significant effect, we computed the 95% confidence intervals (CI) for selection coefficients resulting from the model averaging. A coefficient was significant if its CI non-overlapped with 0. We included these variables to fit average seasonal iSSA models on three different training datasets. Second, we used those averaged seasonal iSSA models to build three different resistance maps for each population. Third, we modeled connectivity between winter and summer ranges using the Randomized Shortest Path algorithm on each resistance map (Panzacchi et al., 2016). Finally, we used 2 different methods to evaluate the performance of our three connectivity models by comparing their predictions to the different validation datasets previously set aside.

#### 1. Sampling training and validation datasets

We built three different combinations of training/validation datasets, which were then used to build resistance maps and model three corresponding connectivity maps. Specifically, we first fitted two seasonal iSSA using 90% of the whole dataset (Figure 2; ‘leave 10% of whole data out’ approach), and second, we fitted population-specific seasonal iSSA models with 90% of the population-specific dataset as a training dataset (‘leave 10% of population data out’ approach). For those two approaches, the remaining 10% of migratory tracks served as validation datasets and the 90%/10% sampling procedure was repeated 250 times to assess uncertainty around connectivity predictions. Third, we fitted iSSA with data from all populations but one, and used data from the discarded population for validation, and repeated this for the 15 populations (‘leave one population out’ approach).

#### 2. Building resistance maps

We then used the different iSSA models to compute seasonal resistance maps that display the relative avoidance of each pixel by an ibex migrating through the landscape. To do so, we multiplied each raster of environmental variables by the corresponding coefficient provided by the iSSA model fitted with a given data source. Then, we summed those rasters and applied the inverse logit function to get habitat selection maps representing the relative probability that an ibex selected a pixel during migration. The RSP algorithm uses a resistance map to model the connectivity, to obtain resistance maps, we applied the inverse function to those habitat selection maps considering that the cost of movement is higher in avoided habitats (Keeley et al., 2016; Zeller et al., 2018).

#### 3. Modeling connectivity between summer and winter ranges

We defined seasonal ranges as 95% kernel areas derived from the corresponding seasonal locations (Worton, 1989), using *kernelUD* function from *adehabitatHR* package (h parameter was set to 400; Calenge, 2022). We restricted our analyses to seasonal ranges connected by migratory tracks to limit our connectivity predictions to the areas actually used by GPS-collared migrant ibex. We used the Randomized Shortest Paths approach to model connectivity (RSP; Saerens et al., 2009; Panzacchi et al., 2016; implemented in the *passage* function from R package *gdistance*; van Etten, 2022) between 10 points randomly sampled within each pair of summer and winter ranges. This algorithm estimates the number of times an ibex would cross each pixel of the resistance map during migration. As in other algorithms relying on the graph theory (e.g., least-cost path and circuit theory), the resistance map is represented as a graph with individuals moving from nodes to nodes (i.e., the center of the pixels) along links/edges with variable costs depending on the values of the resistance map. The RSP computes the least-cost path, the path which minimizes the distance and costs accumulated along a trajectory joining a source and a destination. The RSP algorithm also integrates a stochasticity parameter θ which allows measuring the degree of departure from two extreme strategies, i.e., random-walk (full exploration of neighboring nodes), when θ = 0, or least-cost path (i.e., optimal exploitation of the landscape by minimizing total costs) for the highest value of θ (see Appendix S10). This allows accounting for intermediate strategies between the two most commonly used methods to model movements in connectivity analyses. We obtained two connectivity maps (one per season) for each training dataset using an optimized stochasticity parameter θ (Appendix S10). We finally obtained three unique connectivity maps (one per training dataset) by averaging the two seasonal connectivity maps for each training dataset.

#### 4. Validating connectivity maps

We used two different methods to evaluate the accuracy of our connectivity predictions. First, we ranked each used step traveled by ibex during migration versus the 15 associated available steps they could have travelled (already sampled for iSSA analyses; see section *Habitat selection analyses during migration*) based on connectivity values at the end of each step and assigned them a value between 1 (lowest connectivity) and 16 (highest connectivity; ranking method; McClure et al., 2016; Goicolea et al., 2021). If accurately predicted, the average rank of used steps should be higher than those of available steps. Second, we converted connectivity values to percentile connectivity values (e.g., the 95^th^ percentile corresponds to the 5% highest values of the connectivity map) and delineated five connectivity corridors as the 80^th^, 85^th^, 90^th^, 95^th^, and 99^th^ connectivity percentiles. We then calculated the percentage of ibex locations collected during migration included in each connectivity corridor as a metric of predictive performance of our connectivity models (representation in corridors; Poor et al., 2012; Zeller et al., 2018; Goicolea et al., 2021). As the percentage of ibex locations during migration that fall within a given corridor is strongly dependent on the area of this corridor, we also computed the proportion of locations in the corridor divided by the corridor surface to get an index of accuracy of connectivity predictions (Appendix S11). We applied both validation methods (ranking and representation in corridors) on the three validation datasets we set aside previously to validate our three approaches to compute connectivity maps.

## RESULTS

### Identification of migrant ibex

Among the 1068 seasonal tracks available in our GPS dataset, we identified 337 migratory tracks (169 in spring, 168 in autumn), distributed between multiple winter and summer ranges within each population. On average, the proportion of seasonal tracks identified as migration was 45% (SD 22.5) over the 15 populations, confirming partial migration. However, it varied greatly between populations, from 13% to 75% in the Bargy and Champsaur populations, respectively (considering populations with enough animals to estimate this proportion). On average, migrant ibex traveled 12 km (SD 8) of topographic distance, with population means that varied from 6 to 22 km and an individual maximum of 62 km. In spring, those migratory tracks lasted 3.5 days (SD 3.6) on average and occurred around May 27 (SD 27 days), while in autumn they lasted 6.3 days (SD 6.3) on average and occurred around October 30 (SD 29 days).

### Habitat selection during migration

In both spring and autumn, ibex traveled in areas with less total elevation change (192.4 m on average for 6 hours) than if they had moved randomly (207.1 m, 7% less; significant in 11/15 populations in spring and 13/15 in autumn; Figure 3). They also selected for proximity to refuges from perceived predation risk (slopes > 40°; 11/15 populations in spring and 10/15 in autumn) and avoided north-oriented areas (10/15 populations in both seasons). During autumn migration only, they also avoided areas expected to be the first covered by snow and where snow may accumulate (snow cover index; 7/15). By contrast, neither anthropogenic infrastructures (proximity to ski resorts and roads) nor linear structures considered as potential landmarks (proximity to ridges, valley bottoms and tree lines) influenced ibex migratory tracks during either season.

**Figure 3:**
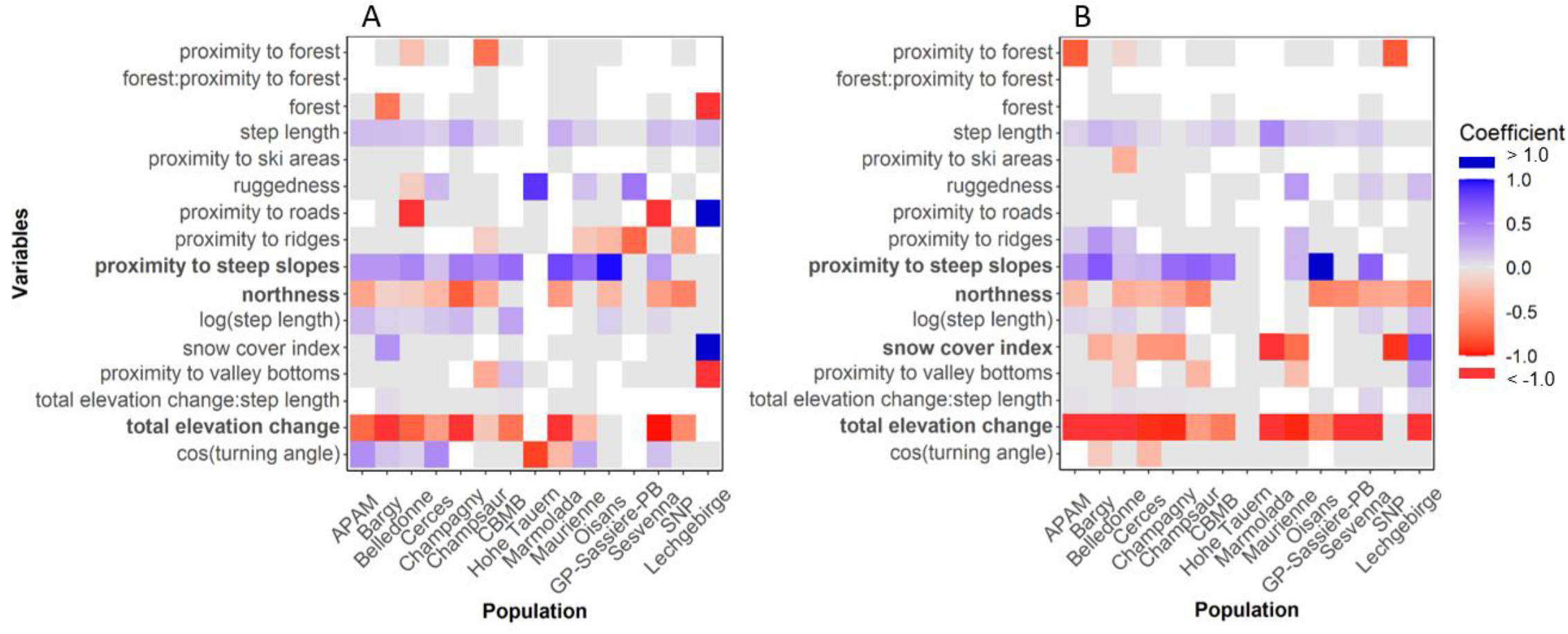
Coefficients provided by population-specific model-averaged (models with ΔAICc < 2) integrated Step Selection Analyses investigating the influence of environmental variables on movement steps performed by Alpine ibex from 15 populations during spring (A) or autumn (B) migration. Blue and red cells represent variables that were selected or avoided for migratory movements, respectively. Grey cells represent non-significant coefficients. We calculated the 95% confidence interval (CI) of coefficients resulting from the model averaging. A coefficient was significant if its CI non-overlaps with zero. White cells represent cases for which the influence of a focal habitat variable could not be tested (not retained during model selection). A variable name is in bold type if its selection coefficient was significant in at least 8/15 populations.

### Connectivity modeling

The three modeling methods performed relatively well and produced similar predictions of ibex migratory corridors. About half of the migratory tracks were in areas with high connectivity, falling in the 95th connectivity percentile corridor (50.4% (SD 14.7) for “leave one population out”; 51.9% (SD 11.8) for “leave 10% of population data out” and 53.1% (SD 13.7) for “leave 10% of whole data out”; Figure 4). The percentage of tracks included in the predicted corridor increased rapidly for lower values of the predicted connectivity corridor, as more than 90% of the tracks were included in the 80th connectivity percentile. The best stochasticity value θ in the Randomized Shortest Path algorithm was equal to 0.1 (Appendix S10). This intermediate value largely outperformed the lower (θ=0; totally random movements) and upper limits (θ=3; deterministic movements) resulting in intermediate connectivity patterns between the diffuse connectivity corridors obtained with the circuit theory approach and the narrow and simple least-cost path that prevented from alternative routes (Figure 5).

**Figure 4:**
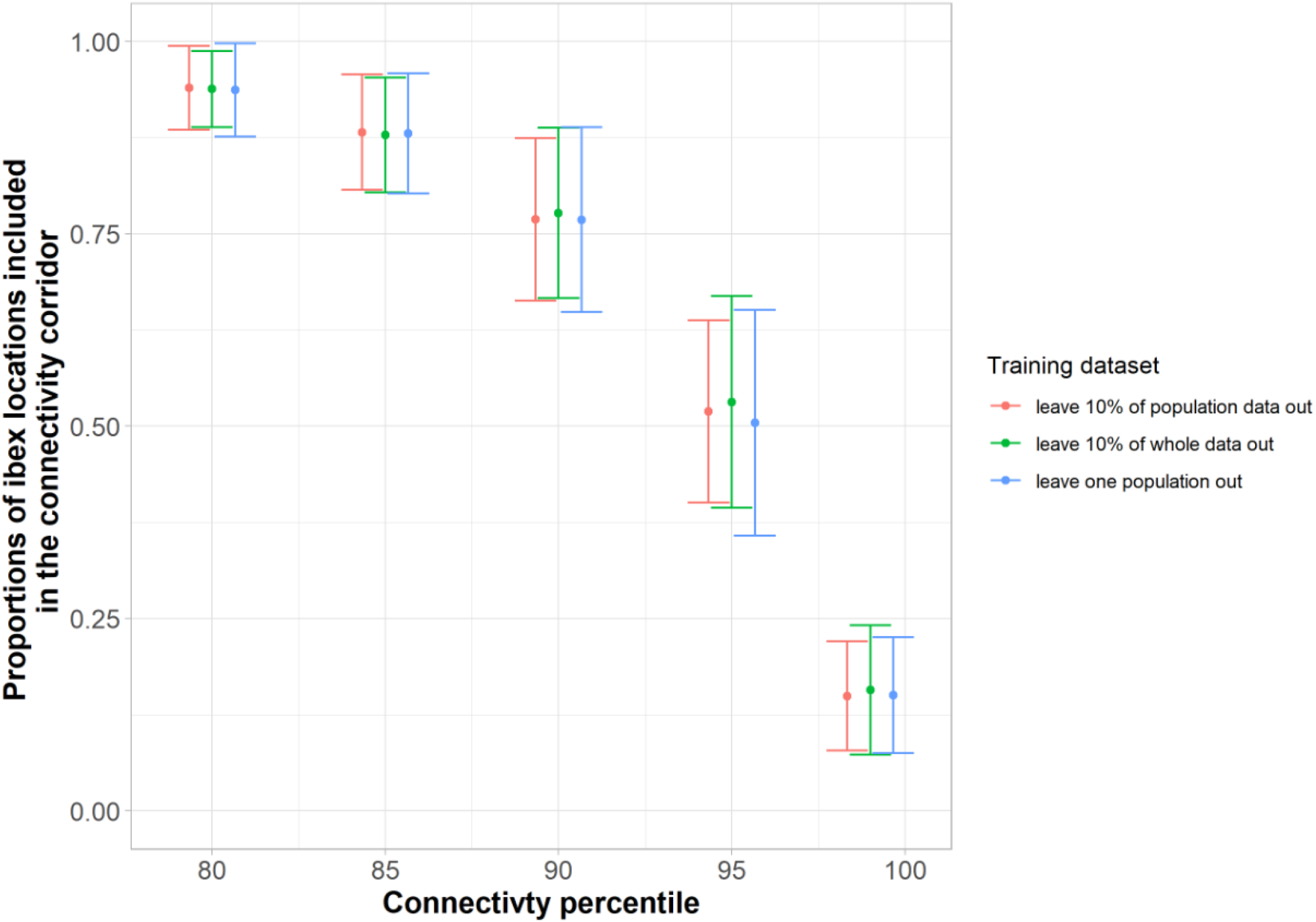
Results of the second validation method, representation in corridors (Goicolea et al., 2021). Proportions of ibex locations from migratory tracks included in the different connectivity corridors defined as the 80th, 85th, 90th, 95th and 99th connectivity percentiles. For the three modeling processes, the mean proportion calculated over the 15 populations is displayed with its standard deviation.

**Figure 5:**
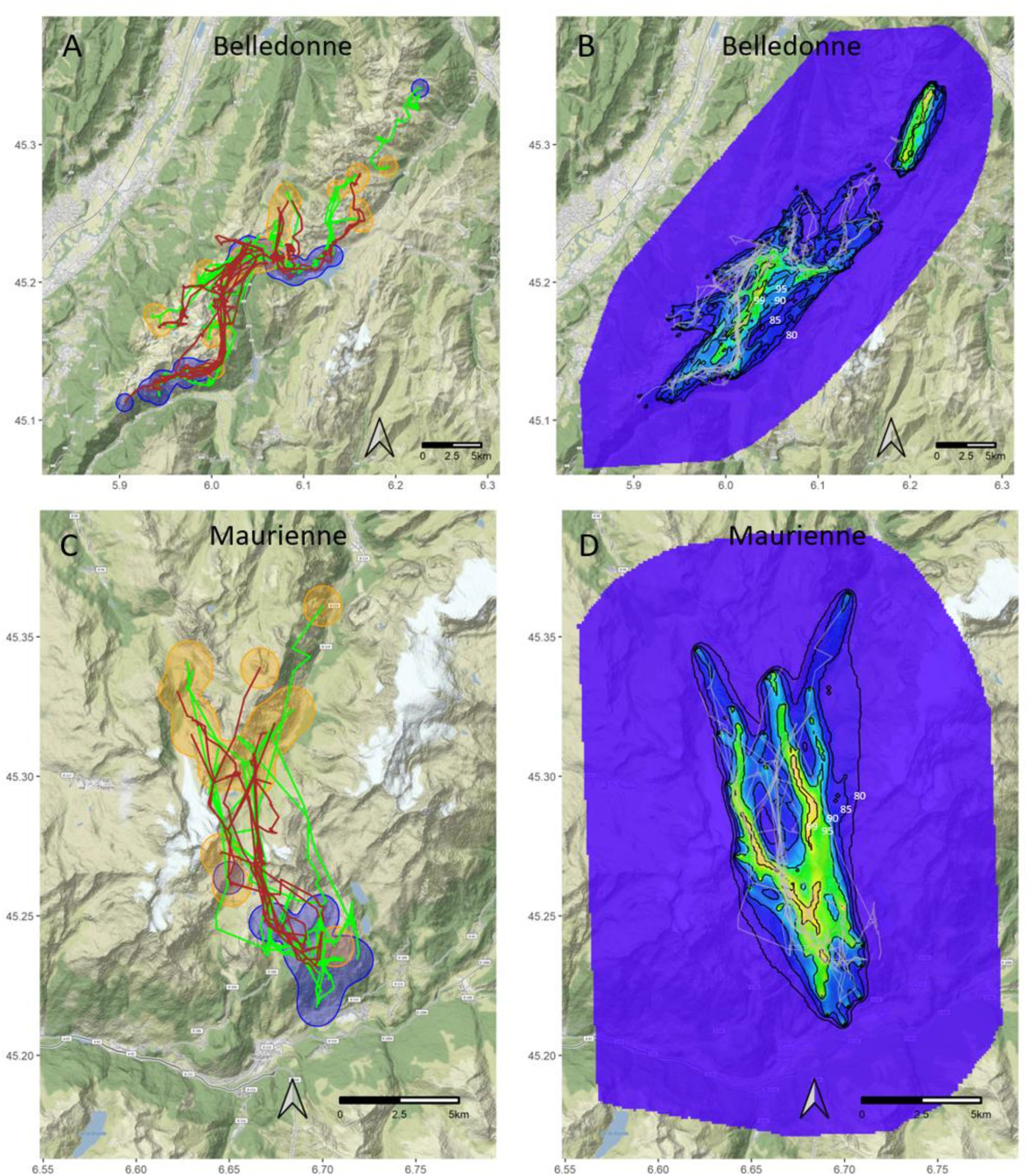
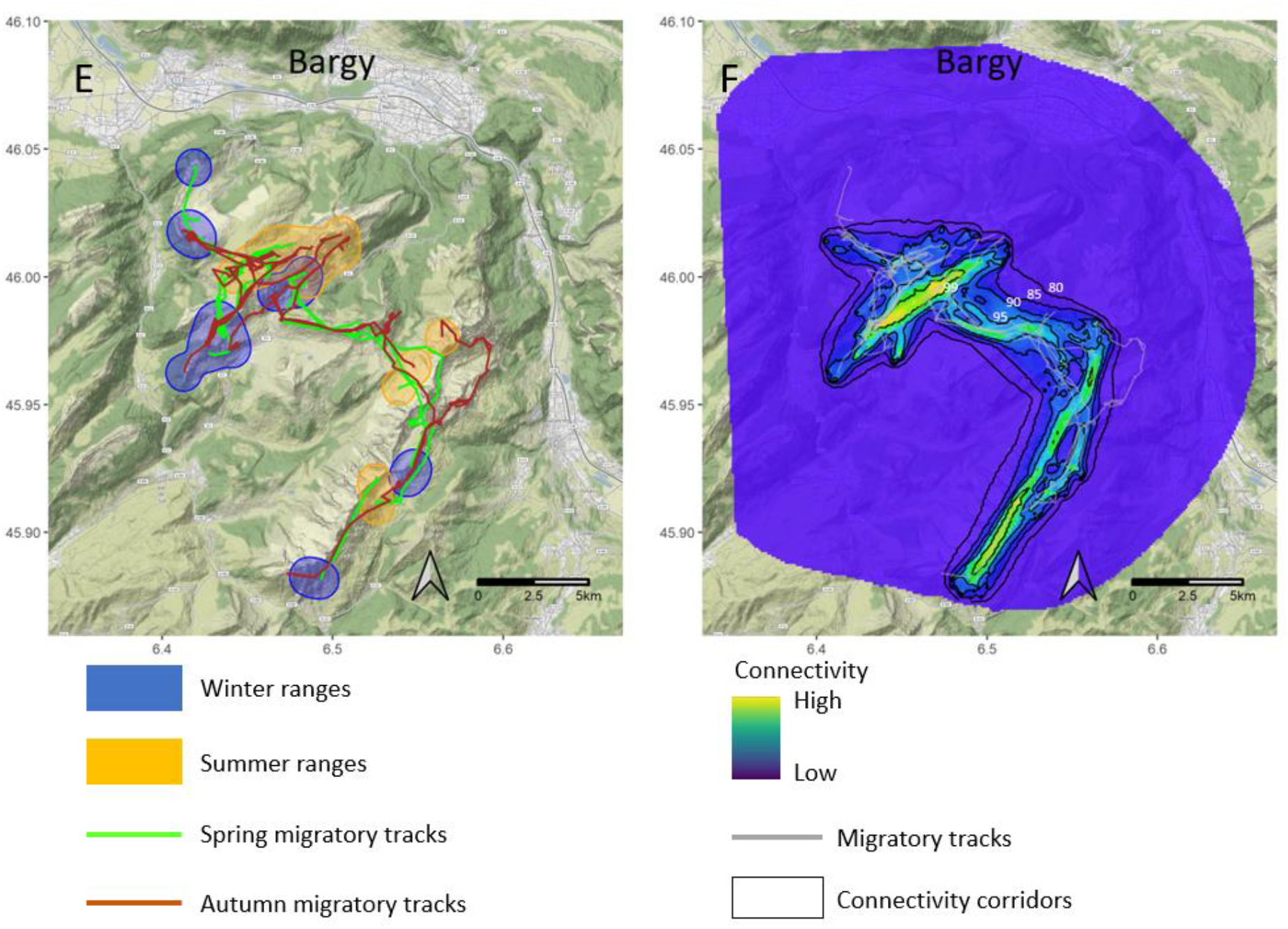
Examples of connectivity modeling. Observed migration routes (spring in green and autumn in brown) and summer and winter ranges (orange and blue) of Alpine ibex from Belledonne, Maurienne and Bargy populations (A, C and E). Connectivity maps obtained from the “leave 10% of whole data out” dataset (B, D and F). The black lines delineate the connectivity corridors as defined in the ‘representation in corridors’ validation method.

According to the ranking validation method, the three connectivity modeling approaches tested (i.e., ‘leave 10% of whole data out’, ‘leave 10% of population data out’ and ‘leave one population out’) provided connectivity maps that predicted ibex migratory movements better than random surfaces (see Figure 5 for examples, and Appendix S12). The mean and median ranks of used steps in the three validation datasets were all>8, with values from 9 to 11 depending on the population, although variability was important (1^st^ quartile and 3^rd^ quartile ranging from 5 to 14 depending on populations; Appendix S13). However, within populations results were similar whatever the training dataset used.

Comparable proportions of locations of migratory tracks were included in connectivity corridors for the three modeling approaches: between 93.7 – 94.0% of locations in the 80th connectivity percentile corridors and between 14.9 – 15.7% in the 99th connectivity percentile corridors (Figure 4, Appendix S14). There was heterogeneity between populations in the accuracy of the predictive models of connectivity, but within the same population, the three connectivity models gave similar results (Appendix S14). The ratio of the proportion of locations included in the corridor over the corridor surface was superior for the 95th and 99th percentile corridors. Therefore, these connectivity corridors captured on average the highest proportion of ibex locations within the smallest surface but the variability in this ratio over the 15 populations was high (Appendix S11).

## DISCUSSION

Relying on a dataset assembling 337 migratory tracks collected in 15 Alpine ibex populations distributed across the Alps, we identified the environmental predictors of corridors in this endemic and emblematic short-distance and altitudinal migrant species. While consistently limiting energetically-costly elevation change, ibex migrated mostly in south-facing snow-free slopes and close to steep areas providing refuges from perceived predation risk. By contrast, neither the landmarks (ridges, tree lines, valley bottoms) hypothesized as visual cues for ibex navigation, nor human infrastructures (ski areas and roads, when present) affected ibex migratory movements. The randomized shortest path algorithm revealed an intermediate movement strategy in Alpine ibex, trading off optimization and exploration during migratory movements. The abilities of the three modeling methods we compared to predict migratory connectivity from the results of those movement analyses, relying either on population-specific or multipopulational approaches, were comparable. They provided efficient connectivity models to inform conservation of Alpine ibex and its habitats, and a framework for future research investigating connectivity in migratory species from multi-populational datasets.

In addition to spring green wave tracking, evidenced in commonly-studied long-distance migrations from North American, Scandinavian and African ungulates, we revealed other predictors, more scarcely investigated, may also drive Alpine ibex during both spring and autumn migration. By focusing on south-facing snow-free slopes, they may partly benefit from emerging vegetation during spring migration (although we did not fully investigate the green wave hypothesis, see Methods; but see Semenzato et al., 2021), while limiting energetically-costly movements in snow-covered areas. Limiting energy expenditures seemed particularly important in Alpine ibex that also strongly avoided high elevation changes during migration, yet relatively short in distance and duration. This behavior may be adaptive in the steep and rugged terrain in which ibex migration occurs (see Passoni et al., 2021 for another example in roe deer *Capreolus capreolus*). Indeed, when traveling through unfamiliar areas for migration, Alpine ibex selected for proximity to steep slopes, i.e. habitats generally used by mountain ungulates to limit perceived predation risk (Grignolio et al., 2007, see also Marchand et al., 2015 for Mediterranean mouflon *Ovis gmelini musimon* × *Ovis* sp.; Baruzzi et al., 2017 in chamois *Rupicapra rupicapra*). Altogether, these results suggest the persistence of the energy- food-cover trade-off, i.e. the most important predictor of ungulate habitat selection all year round (Lima & Dill, 1990; Houston et al., 1993; Mysterud & Østbye, 1999), as a major driver of Alpine ibex migration routes. This trade-off may also explain the intermediate movement strategy of migrant ibex trading off optimization and exploration during migratory movements, as revealed by the randomized shortest paths algorithm. By contrast, none of the landmarks tested seemed to be used by migrant ibex as compass for navigation during migration. Yet, recent studies revealed how natural landscape features can be used by mountain ungulates, including Alpine ibex, to delimit their seasonal home ranges and constitute cognitive maps to gather and memorize spatially explicit information for navigation (Seigle-Ferrand et al., 2022). Further research is hence needed to investigate the importance of other navigation cues/mechanisms, and more generally other drivers of migration corridors identified in other contexts/species (e.g., memory; Bracis & Mueller, 2017, Merkle et al., 2019; social learning and cultural transmission; Jesmer et al., 2018) that were not investigated here in absence of data to do so.

Combined with the randomized shortest paths algorithm, the results of three modeling methods relying either on population-specific or multi-population approaches provided reliable and comparable connectivity maps. Both the “ranking” and “representation in corridors” validation approaches indicated relatively high levels of agreement between connectivity corridors and actual migratory tracks, although the dispersion associated with reliability measurements was high. Indeed, in several populations, some predicted high-use areas were not used by ibex, or inversely, ibex used areas that were not predicted as providing high connectivity. Thus, factors such local idiosyncrasies in landscape features may be involved at the population level.

Despite obvious topographic, climate and anthropogenic differences in the areas used by the 15 studied populations across the whole Alps, contributing to the differences in movement strategies across populations (Figure 3), we chose to fit iSSA with environmental variables that had significant influence in most populations, and to average the resulting resistance maps over both seasons (spring and autumn; see Figure 2). This probably contributed to increasing uncertainty in connectivity predictions as inter-population differences were ignored. However, when assessed with the same validation approach (here “representation in corridors”, easily translatable into management/conservation measures; McClure et al., 2016), the performance of our connectivity models was comparable or better than those reported in other studies (here, 73-78% locations in the 90th percentile corridor, 68-72% in Poor et al., 2012 - pronghorn *Antilocapra americana*; 65% in Zeller et al., 2018 - puma *Puma concolor*; 42% in Goicolea et al., 2021 - Iberian lynx *Lynx pardinus*). Besides, those average connectivity models may be more reliably transferable or generalizable to other populations/contexts over the species’ distributional range and even in the absence of any data. Therefore, it may constitute an invaluable tool for the conservation of this endemic and emblematic species and its habitats.

Even though we did not find any major effects of human infrastructures that could impede ibex migration (i.e. ski areas and roads, probably due to their scarcity in the vicinity of areas where ibex have been reintroduced), climate warming and the development of human activities and infrastructures, particularly present in the Alps (Parmesan & Yohe, 2003; Schmeller et al., 2022), could reshape movement corridors of alpine animals in the near future (see an example with pronghorn, Zeller et al., 2021). In addition, despite the numerical success of the species reintroduction programs over the Alps, Alpine ibex still face important conservation issues (e.g., dramatically low genetic diversity, lack of functional meta- populations; Biebach & Keller, 2009; Brambilla et al., 2021) and migration corridors remain poorly protected outside. In this context, preserving and (re-)establishing connectivity within and between ibex populations will probably be a major conservation issue in the next decades, and tools such as our connectivity models could be particularly helpful.

More generally, our study also provided an original methodological framework for future research and conservation efforts dedicated to connectivity analysis and predictions of movements other than migration. Here, the three different approaches (i.e., ‘leave 10% of whole data out’, ‘leave 10% of population dataset out’ and ‘leave one population out’) revealed no major differences in accuracy of corresponding connectivity predictions. Thus, our models could be used to predict migratory movements in monitored populations with either enough data, using population-specific model, or using data from all populations. Moreover, we could predict movements in populations where no GPS data are available but seasonal range locations are known or predicted with habitat selection models. With the advent of animal tracking over the last decades, and the generalization of initiatives aiming at gathering those GPS data in common databases (e.g. Movebank, Euromammals, Biologging initiative, Global Initiative for Ungulate Migrations; Kauffman et al., 2021; Urbano, Cagnacci & Euromammals consortium, 2021), multi-population analyses will develop and testing the reliability of population-specific versus multi-population connectivity predictions is crucial, particularly in a context of demand and need around conserving and restoring connectivity within species distribution ranges.

## ACKNOWLEDGMENTS

We thank all the professionals and interns involved in the monitoring of GPS-collared ibex on all study sites, in particular, F.Couilloud and le Service Départemental de la Haute-Savoie et le Service Départemental de l’Isère from the Office Français de la Biodiveristé, and L.Gautero, V.Roggero, A.Rivelli, E.Piacenza, M.Dotto, A.Menzano. We thank the Alcotra LEMED-ibex and all its partners: Ecrins NP, Gran Paradiso NP, Asters-CEN 74, Vanoise NP, Autonomous Region Aosta Valley, Mercantour NP, Protected areas of Maritime Alps, Protected areas of Cottian Alps – for providing GPS data collected during the program.

We also thank the GSE-AIESG: Gruppo Stambecco Europa-Alpine Ibex European Specialist Group and Gran Paradiso National Park for providing data on ibex population distribution, and in particular A.Brambilla. We warmly thank the Global Initiative for Ungulate migrations and its mapping team, the OFB, ANR MovIt, ANR HUMANI, CEFE and LECA teams for fruitful discussion on early versions of this manuscript.

This research and the data collection were founded by the Office Français de la Biodiversité, Agence Nationale de la Recherche (ANR; Grant/award Numbers: HUMANI #18-CE03-0009, Mov-It #16-CE02-0010), the Alcotra LEMED-ibex. The Stiegl Brewery of Salzburg for the Hohe Tauern National Park. The Swiss National Park. The University of Padova, project n. CPDA094513/09 MIUR ex 60%, projects 60A08-2154/14 and 60A08-2017/15; Veneto region, Wildlife and Hunting Service (Regione Veneto-Unità di Progetto Caccia e Pesca); Fondazione Edmund Mach, ordinary funds from Autonomous Province of Trento. The Research Council of Norway, project nr 223257. Mercantour National Park was financed by GMF. The study in Vorarlberg (Austria) was funded by a hunting community (private financing).

