## Supporting information for "Identifying the environmental drivers of corridors and predicting connectivity between seasonal ranges in multiple populations of Alpine ibex (*Capra ibex*) as tools for conserving migration"

Appendix S1: Details on the studied Alpine ibex *Capra ibex* populations and GPS monitoring

| Country | Population name/ (massif) | Longitude/Latitude  Mean altitude | Population ID and Name in Brambilla et al. 2020 | Monitoring period | Number of ibex tracked (number of potential migrations available) | GPS schedule (number of GPS locations/time units) |
| --- | --- | --- | --- | --- | --- | --- |
| Austria | Hohe Tauern | 12.8°E/47.1°N  2400 m | AUSA02  Hohe-Tauern | 2006-2008; 2010-2011; 2017-2018 | 7 (13) | 1 loc/3 h  and  1 loc/4 h |
|  | Lechgebirge | 10.0°E/47.2°N  2000 m | AUVO03  AUVO05  Kleinwalsertal-Klostertal  Alberg Valluga | 2007-2010 | 12 (22) | 1 loc/3 h |
| France | Bargy | 6.5°E/46°N  1900 m | FRV08 Bargy | 2013-2020 | 117 (270) | 1 loc/1 h |
|  | Belledonne | 6.1°E/45.2°N  2100 m | FRV12 Belledonne | 2017-2020 | 35 (86) | 1 loc/1 h and 1 loc/2 h |
|  | Champagny | 6.8°E/45.5°N  2300 m | FRV06 Champagny-Peisey | 2018-2020 | 13 (24) | 1 loc/1 h and  1 loc/3 h |
|  | Maurienne | 6.7°E/45.3°N  2500 m | FRV01 Maurienne | 2018-2020 | 12 (28) | 1 loc/3 h and  1 loc/6 h |
|  | Contamines – Beaufortain – Mont Blanc  (CBMB) | 6.7°E/45.7°N  2200 m | FRV09 Contamines - Beaufortain - Mont-Blanc | 2018-2020 | 11 (46) | 1 loc/3 h and  1 loc/6 h |
|  | Oisans | 6.0°E/44.9°N  2200 m | FRV15 Valbonnais-Oisans | 2013-2020 | 20 (46) | 1 loc/3 h and  1 loc/6 h |
|  | Champsaur | 6.2°E/44.8°N  2300 m | FRU03 Vieux-Chaillol-Sirac | 2013-2020 | 43 (120) | 1 loc/3 h and  1 loc/6 h |
|  | Cerces | 6.5°E/45.1°N  2400 m | FRU02 Cerces-Galibier | 2015-2020 | 22 (64) | 1 loc/3 h and  1 loc/6 h |
| France - Italy | Alpi Marittime (Italy) - Mercantour (France)  (APAM) | 7.1°E/44.2°N  2300 m | FRU10  FRU01  ITCN01  Nord-ouest Mercantour  Est Mercantour  Alpi Marittime | 2018-2020 | 36 (126) | 1 loc/3 h and  1 loc/6 h |
|  | Grand Paradiso National Park (Italy) – Sassière (France) – Prariond Bonneval (France)  (GP-Sassière-PB) | 7.1°E/45.5°N  2600 m | FRV14  FRV05  ITTO05  Sassière-Prariond  Bonneval-sur-Arc  PNGP-Valle dell’Orco | 2003-2004 | 19 (32) | 1 loc/3 h |
| Italy | Marmolada | 11.9°E/46.4°N  2300 m | ITTN04 Monzoni-Marmolada | 2010-2018 | 32 (56) | 1 loc/1 h |
|  | Sesvenna | 10.4°E/46.7°N  2600 m | ITBZ03  CHGR05  Sesvenna  Macun-Terza-Sesvenna | 2018-2020 | 13 (30) | 1 loc/1 h |
| Switzerland | Swiss National Park (SNP) | 10.1°E/46.6°N  2400 m | CHGR02  Albris | 2007-2019 | 30 (102) | 1 loc/1 h  1 loc/2 h  and  1 loc/4 h |

Appendix S2: Parameters used in Migration Mapper

GENERAL VARIABLES

- processing cores: by default

- maximum speed: 10 km/h

- mortality parameters: * minimum distance: 100 m (# death = 5 days within a circle of 100 m) * time unit = 120 hours

- displacement type: Displacement

- use parallel processing: Yes, use parallel processing

Brownian Bridge Processing Variables

- BM Var: 0

- Time step: 5

-Mult 4 Buff: 0.2

- Minimum Area: 0

- Raster resolution: 25 m

- Corridor percentile: 99

- Simplify: checked/TRUE

- Tolerance: 25 m

- Stopover percents: 10

- Corridor percents: * Corridor percents medium: 10 * Corridor percents high: 20

- BB Type: normal

- Information criteria: LOOCV

- Maximum Fix Interval (hours): 24 hours

- Location error: 20

VARIABLES SPECIFIC TO WINTER RANGE ANALYSIS

- Use same annual dates: False/Unchecked

- Core contours - Winter: By default

- Minimum Days: 10

- Quantile Selectors – Winter * quartile end falls: 0.95 * quartile Start Spring: 0.5

Appendix S3: Example of determination of tracks considered as seasonal migration and migration periods in the Migration Mapper application.
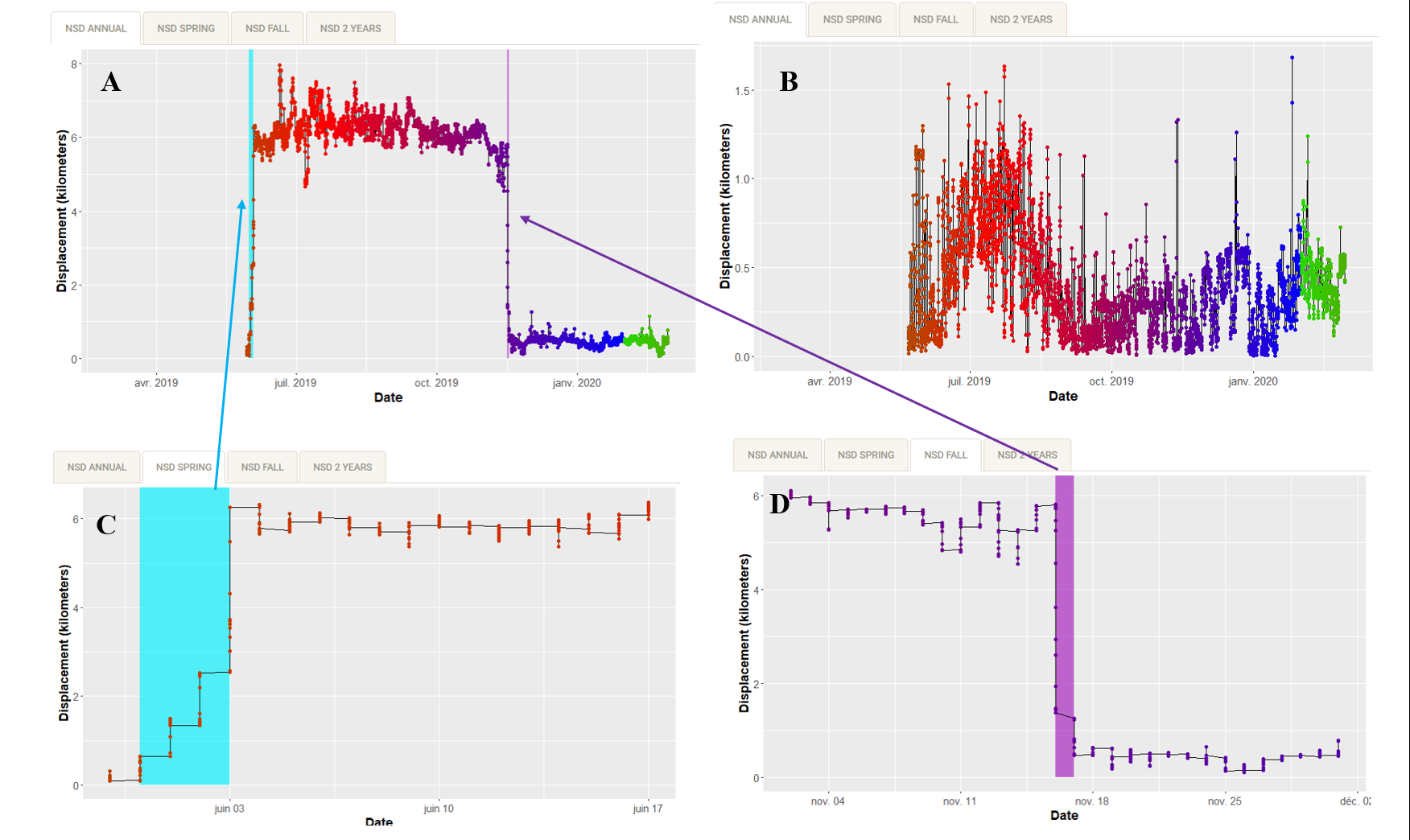

Two examples of annual variation in Net Squared Displacement (NSD): (A) in an Alpine ibex considered as migrant both in spring and autumn, (B) in an ibex considered as resident. Identification of spring (C) and fall (D) migration periods and locations. We identified these periods as the last location preceding the increase/decrease in the NSD and the first location when the NSD stabilizes.

Appendix S4: Planimetric distance and altitudinal interval between seasonal ranges and overlap of seasonal ranges of migrant and resident Alpine ibex

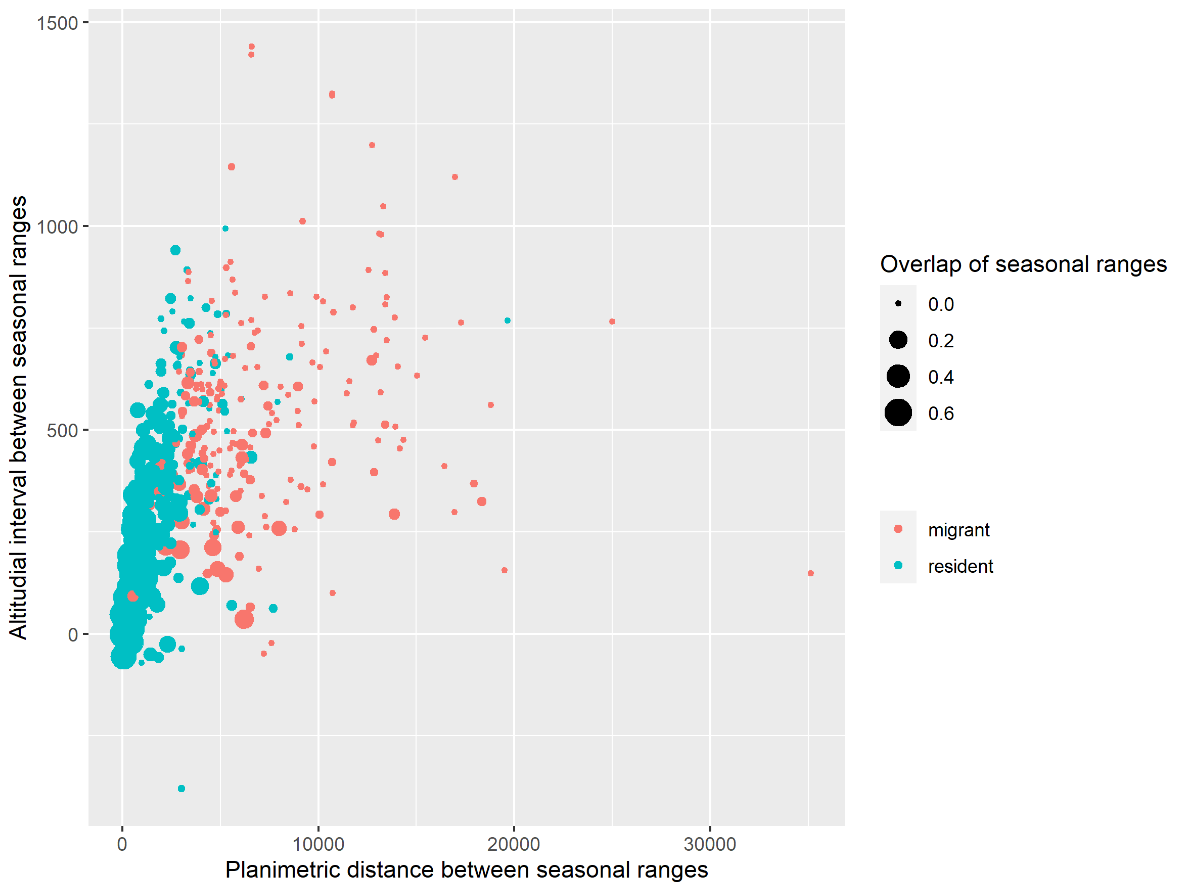

The distances were calculated between the centroid of seasonal ranges of Alpine ibex. Seasonal ranges were defined as the contour 95 of the kernel density estimated with the locations from July 15^th^ to September 1^st^ for summer and November 30^th^ to March 1^st^ for winter ranges. We calculated the overlap of seasonal ranges as the volume of intersection between the two seasonal ranges. Mean overlap of seasonal ranges for migrants: 0.02 (±0.04); Mean overlap for residents: 0.13 (±0.16); Mean distance between seasonal ranges for migrants: 7236 m (±4613); Mean distance between seasonal ranges for residents: 2084 m (±1831); Mean altitudinal difference between seasonal ranges for migrants: 546 m (±259); Mean altitudinal difference between seasonal ranges for residents: 346 m (±217).

Appendix S5: Sources and description of environmental covariates

| Variable | Description | Unit / Range of values | Resolution | Source |
| --- | --- | --- | --- | --- |
| Total elevation change | Sum of changes in elevation values of each grid cell crossed along a step | meters | 25m | Calculated with European Digital Elevation Model (EU-DEM), version 1.1  <https://land.copernicus.eu/imagery-in-situ/eu-dem/eu-dem-v1.1> |
| Vector Ruggedness Measure | Ruggedness measure quantifying the dispersion of vectors orthogonal to the terrain surface | Unitless | 25m | Sappington et al. 2007  *vrm* function from package *spatialEco* in R applied on the European Digital Elevation Model (EU-DEM), version 1.1  <https://land.copernicus.eu/imagery-in-situ/eu-dem/eu-dem-v1.1> |
| Northness | Cosine of aspect derived from the DEM | From -1 to 1 when orientation goes from south to north | 25m | *terrain* function from package *raster* in R applied on the  Elevation Model (EU-DEM), version 1.1  <https://land.copernicus.eu/imagery-in-situ/eu-dem/eu-dem-v1.1> |
| Snow cover index | calculated as the total annual number of days a pixel was covered by snow | days | Spatial resolution: 500 m  Temporal resolution: 8 days | Hall & Riggs 2021  MODIS Snow [product MOD10A2], NASA  <https://modis-snow-ice.gsfc.nasa.gov/?c=MOD10A2> |
| Forest | Presence of forest cover | 0/1 (Absent/Present) | 25m | Corine Land Cover 2012  <https://land.copernicus.eu/pan-european/corine-land-cover/clc-2012?tab=download> |
| Proximity to steep slopes (>40°) | Negative exponential function applied to the distance to the nearest slope >40°  The threshold used for the exponential negative function to compute proximity is set to 500m (beyond 500m proximity equals 0) | 0 to 1 when proximity increases | 25m | Based on slope raster derived from the EU-DEM |
| Proximity to ridges | Negative exponential function applied to the distance to the nearest ridge  The threshold used is 500m | 0 to 1 when proximity increases | 25m | Derived from the EU-DEM using the r.param.scale tool in GRASS GIS 6.4.  4 (Neteler et al. 2012)  We took a minimum size of 500 ha to delimit the watershed (25m*25m*8000 cells) |
| Proximity to valley bottoms | Negative exponential function applied to the distance to the nearest valley  The threshold used is 500m | 0 to 1 when proximity increases | 25m | Derived from the EU-DEM using the r.param.scale tool in GRASS GIS 6.4.  4 (Neteler et al. 2012)  We took a minimum size of 500 ha to delimit the watershed (25m*25m*8000 cells) |
| Proximity to tree lines | Negative exponential function applied to the distance to the nearest tree line  The threshold used is 500m | 0 to 1 when proximity increases | 25m | Derived from the forest cover (Corine Land Cover) |
| Proximity to roads | Negative exponential function applied to the distance to the nearest road  The threshold used is 500m | 0 to 1 when proximity increases | 25m | Derived from roads of OpenStreetMap  Using *osmdata* package in R  <https://cran.r-project.org/web/packages/osmdata/vignettes/osmdata.html>  Download:  2021/09/20 |
| Proximity to ski areas | Negative exponential function applied to the distance to the nearest ski area  The threshold used is 500m | 0 to 1 when proximity increases | 25m | Derived from OpenSnowMap data  <https://www.opensnowmap.org/iframes/data.html>  Download: 2020/12/04. |

Appendix S6: Description of Used Habitat Calibration plots (UHC plots)

UHC plots allow checking the predictions of mixed-effect Poisson models (Fieberg et al. 2018). Such a model is well calibrated if the distribution of observed values of used steps (black line) falls within the distribution of predicted values (grey). The values at available steps are represented with a red dotted line. This method consists in fitting models with a subset of data (here 90% of the dataset) and then sampling beta coefficients from normal distributions N(β, SE(β)²). This sampling is realized 1000 times and the beta coefficients are used to fit models on the remaining 10% of the data. Then, we can visually compare the distributions of values at used steps predicted by the models and the actual (observed) values at these used steps. This comparison allows to infer whether the models correctly predict ibex habitat selection.

UHC plots were first used to check if the models were correctly calibrated. To do so, we used the mixed-effects Poisson models built as described in Appendix 6. We made predictions on a validation dataset consisting of 10% of the randomly sampled data that we set aside (A for spring model and B for autumn model). The UHC plots were then used to assess the effect of sampling intensity on model reliability: to do so, we made predictions using models fitted with 1 and 2h steps on data with 6h steps (C in spring and D in autumn).

The UHC plots revealed for most variables that the distributions of used step values fall within the distributions of values predicted by our average models (Poisson models; Appendix 6). When fitting the models with 1 or 2h steps and predicting values for 6h steps, we also observed that there were no major differences in habitat selection patterns for most variables.

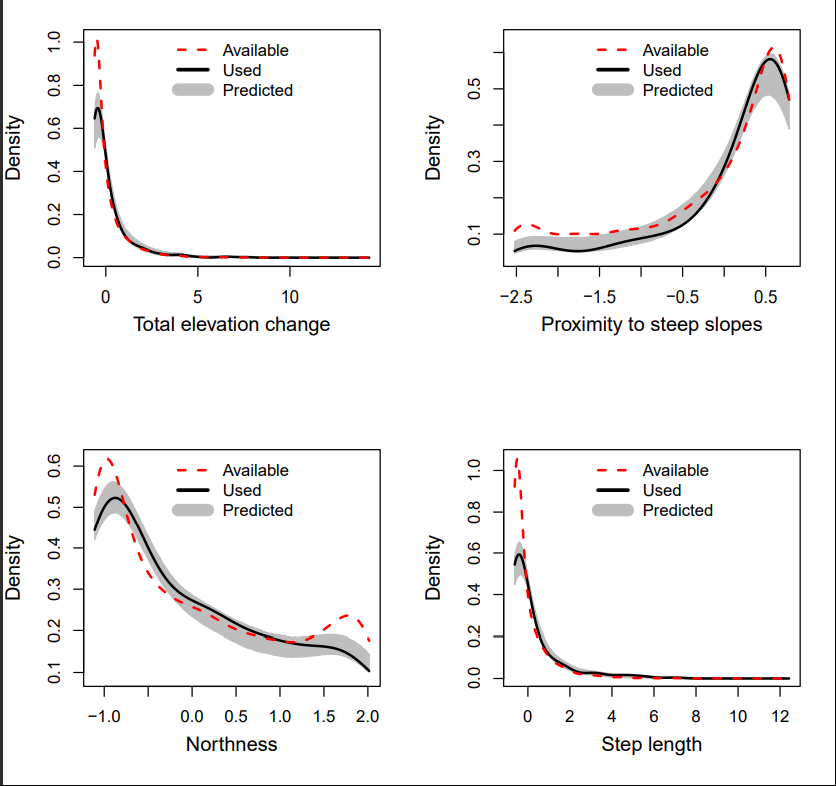

A

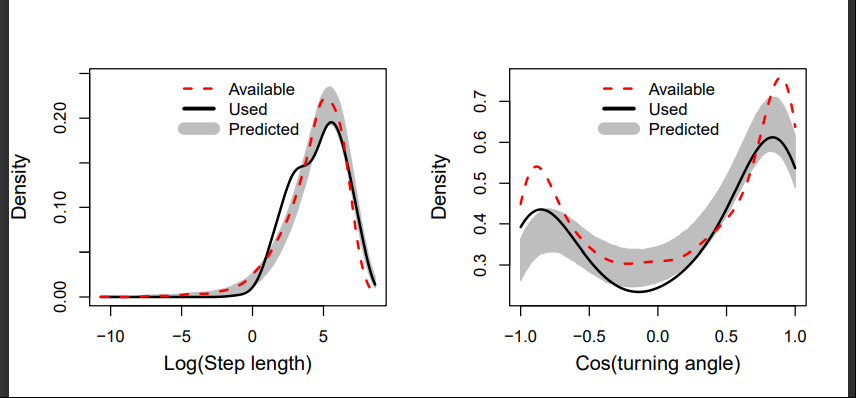

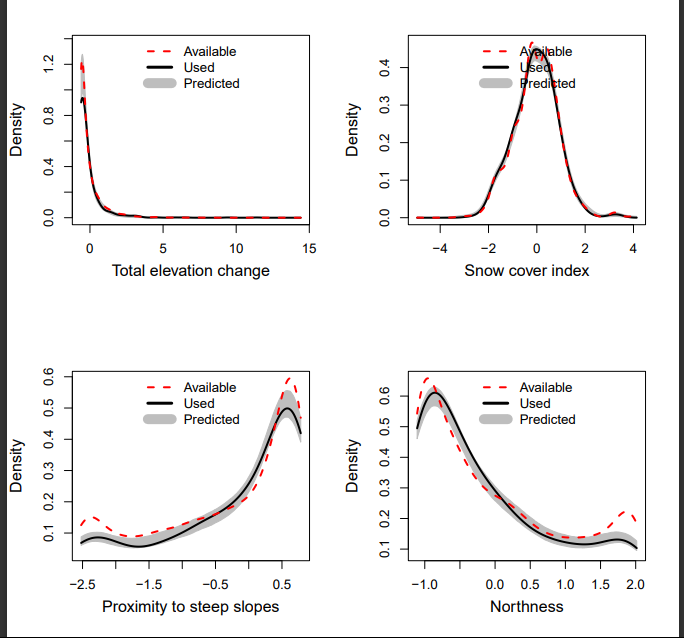

B

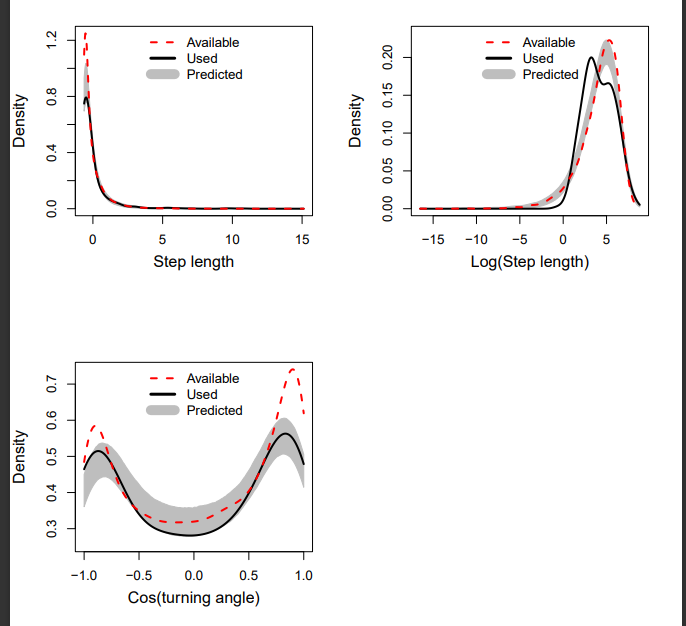

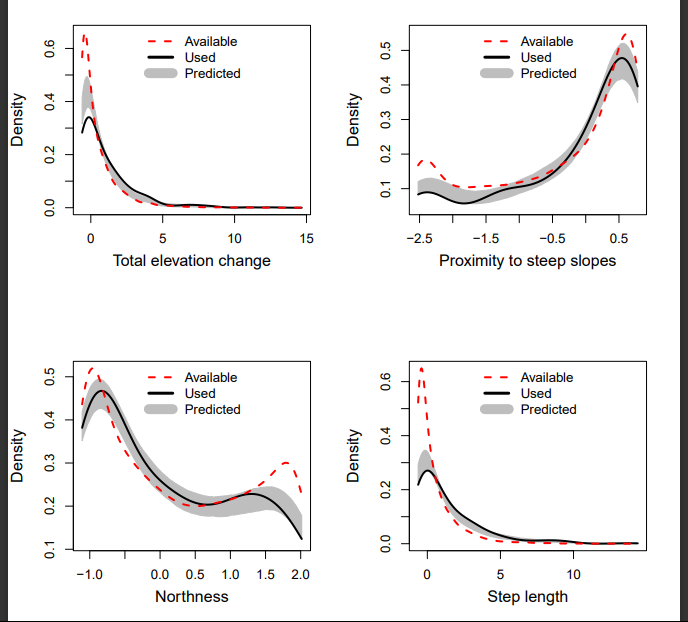

C

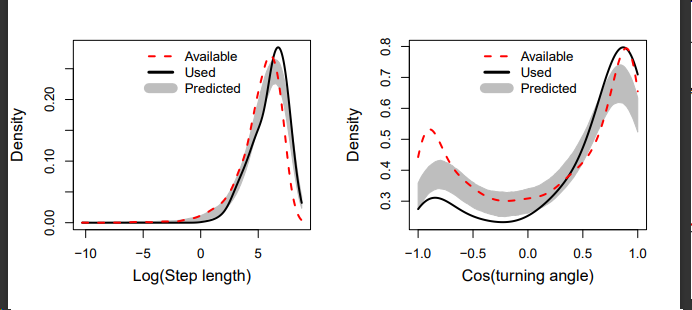

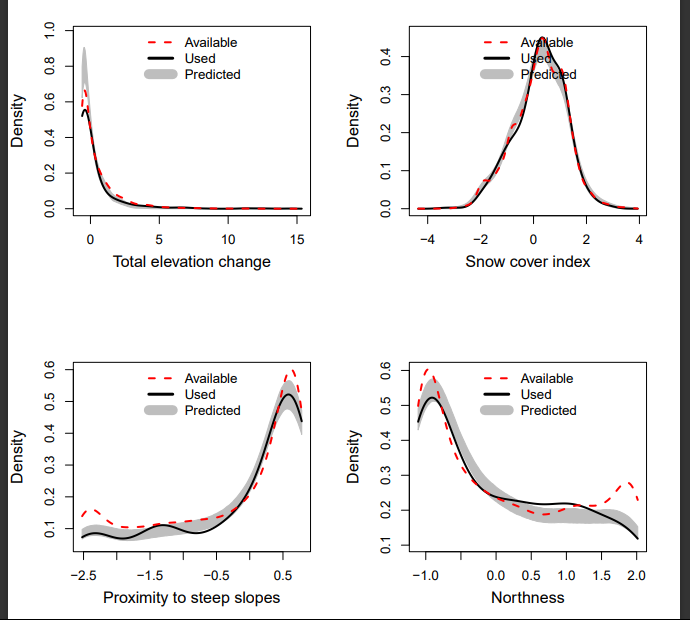

D

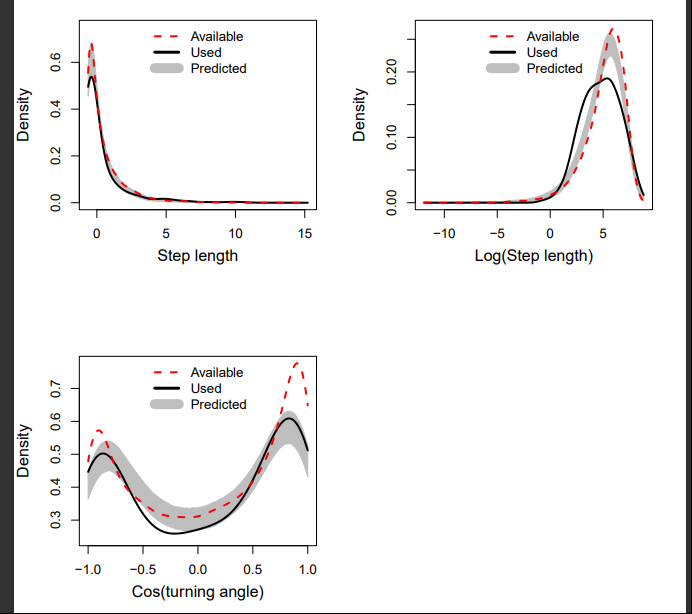

Used Habitat Calibration plots showing the agreement between model predictions and used steps. The UHC plots were made using 90% of the dataset to fit the model and 10% to make the predictions (A in spring and B in autumn). Then, we fitted the model with the data corresponding to the 1h and 2h time steps and made the predictions over the 5h and 6h time step data (C in spring and D in autumn). Grey areas represent the 95% confidence intervals for the predictions, black lines represent the values at and along used steps and red dotted lines represent values at or along available steps.

Appendix S7: Number and sex of migrant Alpine ibex (individual-years, identified as migrant if migrated at least once during the year of monitoring)

| **Population** | **Male migrants** | **Female migrants** | **Migrants with unknown sex** | **Total migrants** |
| --- | --- | --- | --- | --- |
| APAM | 31 | 20 | 17 | 68 |
| Bargy | 18 | 0 | 0 | 18 |
| Belledonne | 27 | 2 | 0 | 29 |
| CBMB | 16 | 0 | 0 | 16 |
| Cerces | 12 | 5 | 0 | 17 |
| Champagny | 3 | 2 | 0 | 5 |
| Champsaur | 25 | 20 | 0 | 45 |
| GP-Sassière-PB | 6 | 1 | 0 | 7 |
| Hohe Tauern | 3 | 3 | 3 | 9 |
| Marmolada | 0 | 3 | 0 | 3 |
| Maurienne | 6 | 6 | 0 | 12 |
| Oisans | 12 | 4 | 0 | 16 |
| Sesvenna | 6 | 3 | 0 | 9 |
| SNP | 8 | 3 | 0 | 11 |
| Lechgebirge | 5 | 0 | 0 | 5 |

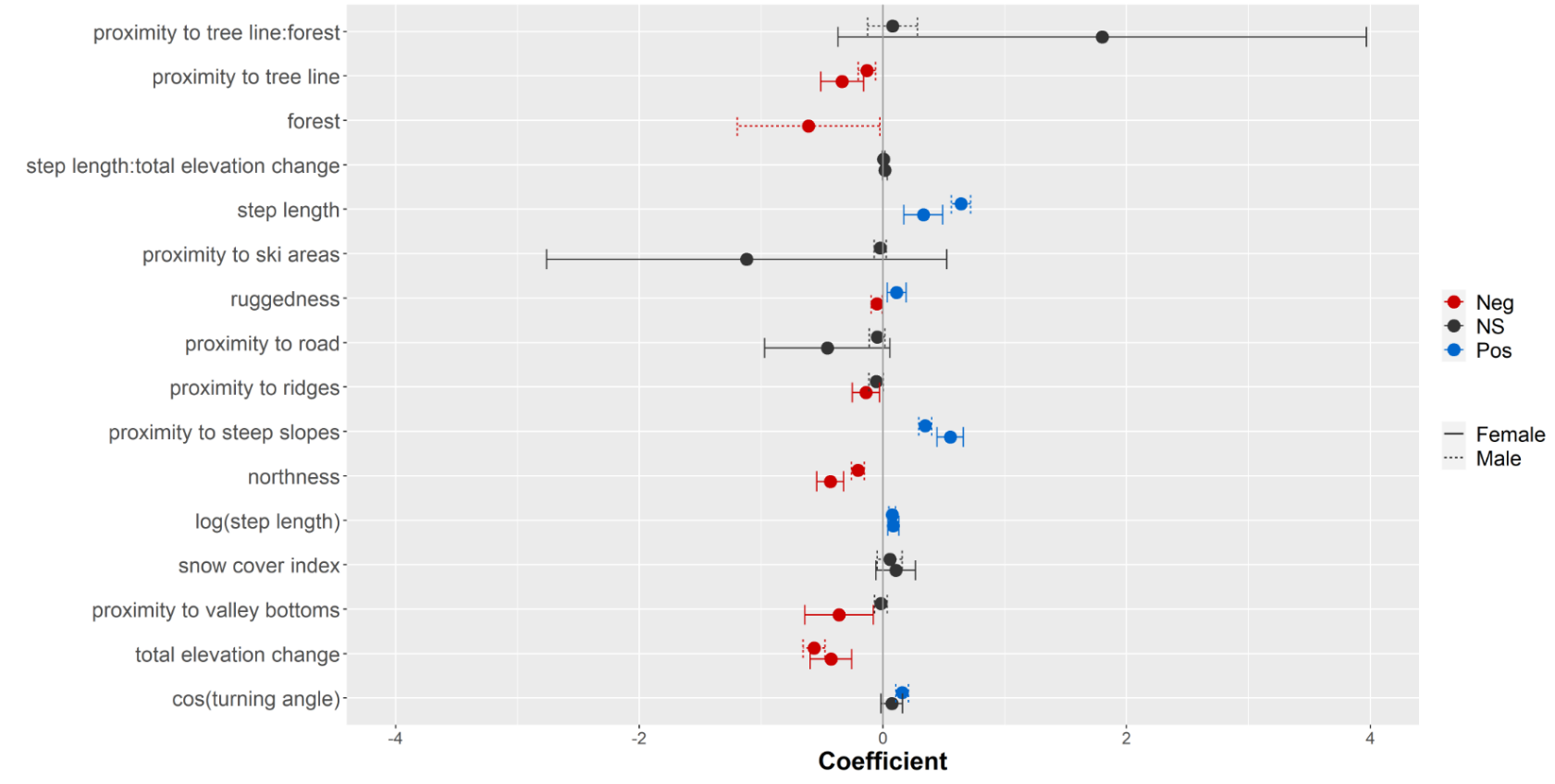

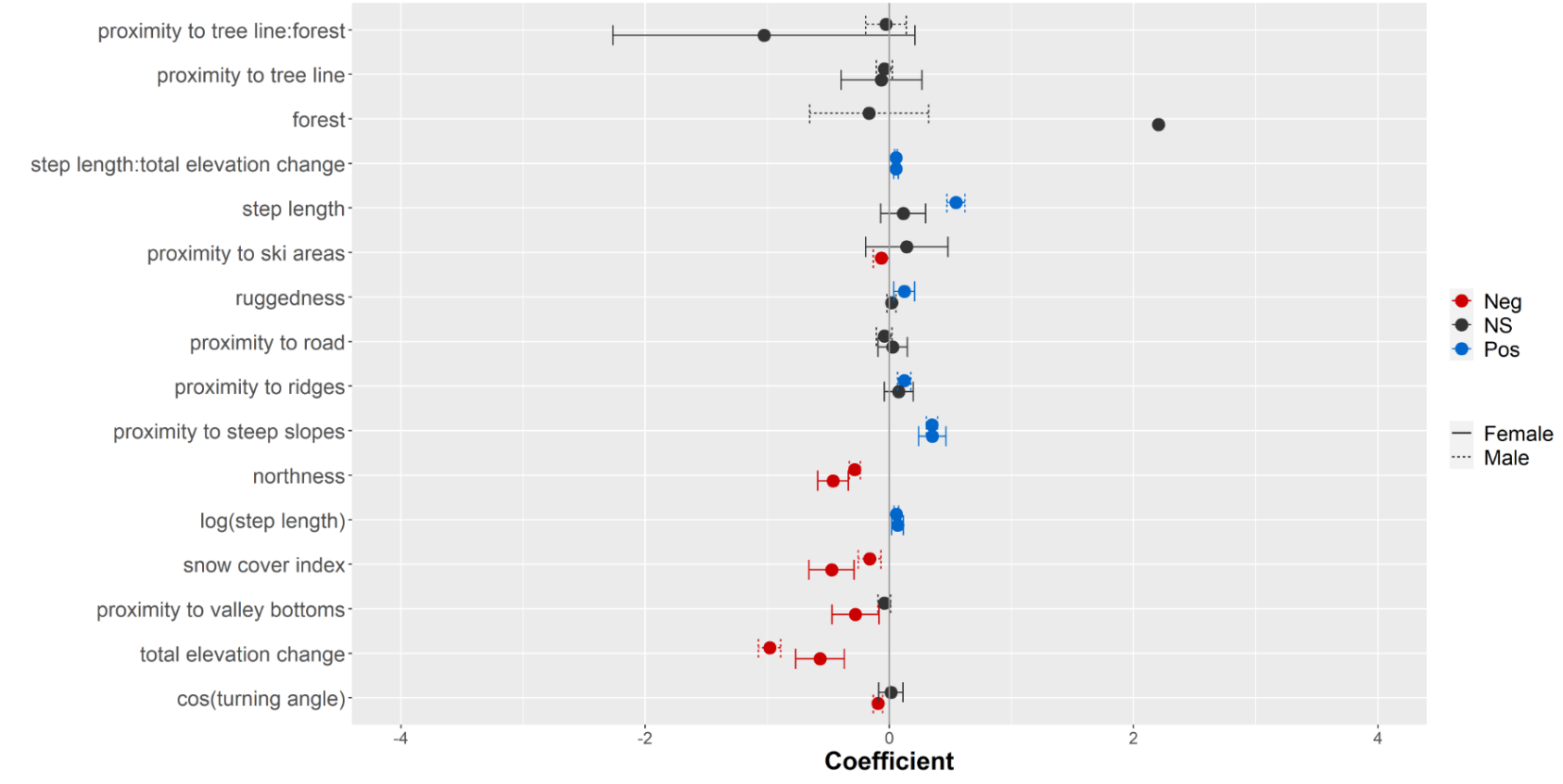
Appendix S8: Sex-specific differences in habitat selection during migration

**B**

**A**

Coefficients from the integrated step selection analyses fitted with data from male (dashed line) and female (solid line) migrants in spring (A) and autumn (B). Variables with a significant positive coefficient are displayed in blue, variables with significant negative coefficient are in red, non-significant coefficients are in gray. To better visualize the results, confidence intervals and/or coefficients of the variable forest were removed (in spring, selection coefficient for forest and females: -6.1[-13.25:0.97]; in fall, selection coefficient for forest and females: 2.20[-0.89:5.30]).

Appendix S9: Coefficients of the mixed-effects Poisson models fitted with data from all populations and the variables with a significant effect in at least 8 populations for spring and autumn migrations. Models were used in the Used Habitat Calibration plots procedure.

We fitted two average models (one per season) on data from all populations. We included in both models the environmental variables that were significant in at least 8/15 populational and seasonal models. Those average models included the variable individual as a random effect to account for repeated measurements on the same individual. We used the method described in Muff et al. (2020; *glmmTMB* function from the package *glmmTMB*) to fit mixed-effect Poisson models. These models allowed us to estimate the mean effects of each variable for the 15 populations and were used in the Used Calibration plots procedure (see Appendix 5).

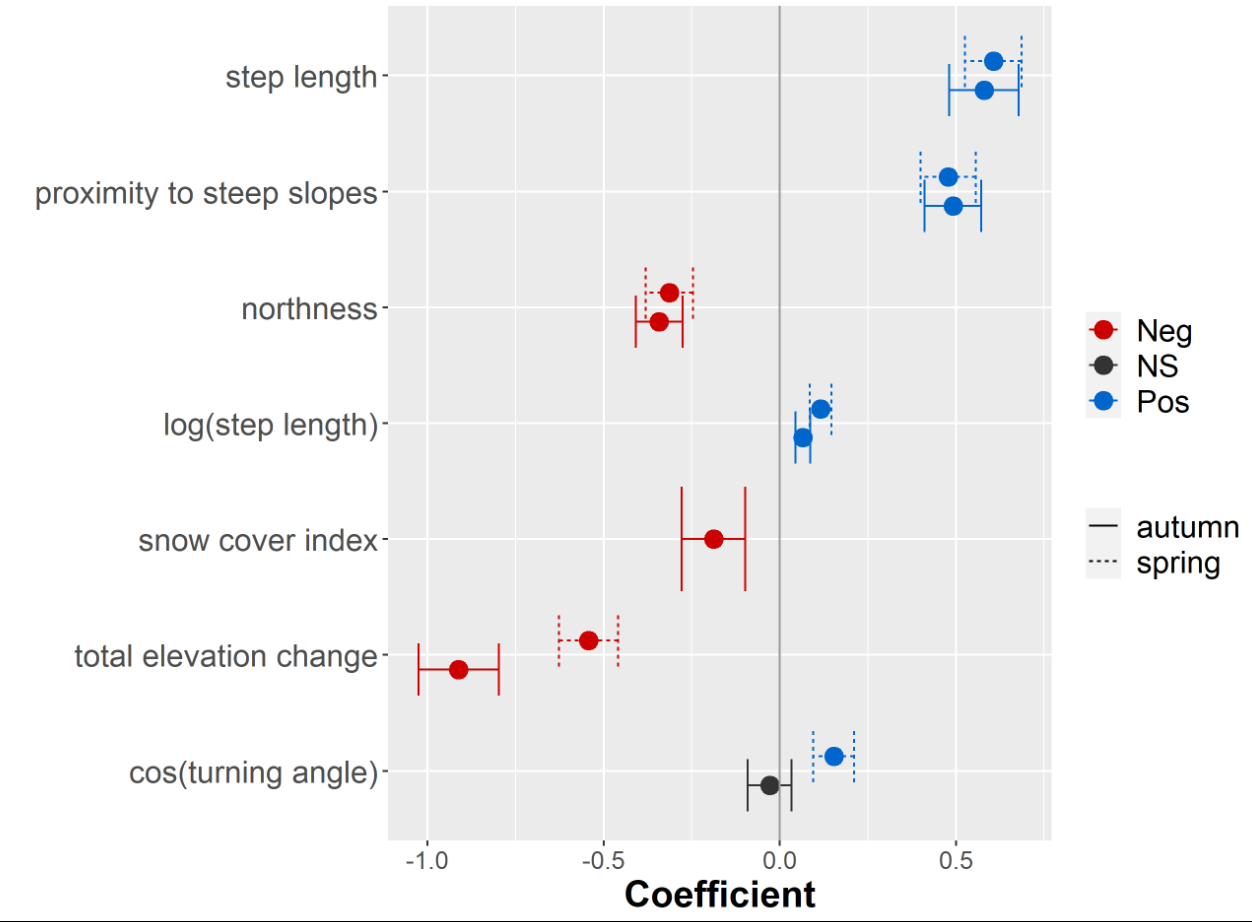
The results were consistent with populational results (results of populational iSSAs, Figure 3). The variables with a significant effect on ibex migratory movements were, by order of importance, total elevation change, proximity to steep slopes, and northness in spring, and total elevation change, proximity to steep slopes, northness and snow cover index in autumn.

Appendix S10: Optimization of the stochasticity parameter in the RSP algorithm

To identify the best stochasticity value *θ* in the Randomized Shortest Path algorithm among 7 different values (0, 0.01, 0.005, 0.1, 0.5, 1, 3; with increasing values corresponding to more deterministic movements), we first derived the corresponding connectivity surfaces from the RSP algorithm and a resistance map derived from the iSSA model averaged over the 15 populations and both seasonal migrations (spring and autumn) (same models as in the “leave 10% of whole data out” procedure). Then, using movement steps (used and available steps) from the validation dataset of the “leave 10% of whole data out” procedure, we fitted seven iSSA (one per θ value / connectivity surface) modeling selection of connectivity surfaces by ibex. We considered the best θ value to be the one for which the AICc of the corresponding iSSA model was the lowest (following Goicolea et al. 2021).

AICc values of the iSSA models fitted to investigate the influence of the stochasticity parameter θ from the Random Shortest Path algorithm on its ability to accurately model the selection of the corresponding connectivity surfaces by Alpine ibex *Capra ibex*.

| θ | AICc | ∆AICc |
| --- | --- | --- |
| 0 | 81817 | 164 |
| 0.01 | 81756 | 103 |
| 0.05 | 81697 | 44 |
| **0.1** | **81653** | **0** |
| 0.5 | 81708 | 55 |
| 1 | 81711 | 58 |
| 3 | 81851 | 198 |

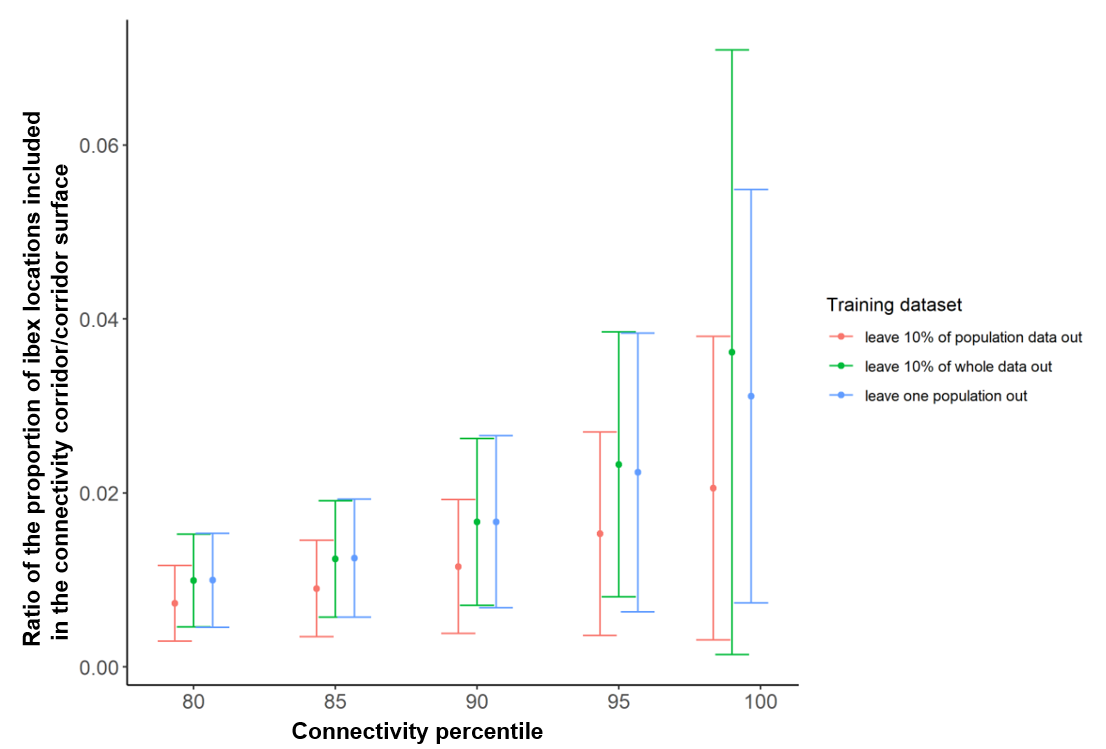
Appendix S11: Extension of the representation in corridors validation method: Ratio of the proportion of locations over the corridor surface.

Mean and standard error, over the 15 populations, of the ratio of the proportion of ibex locations from migratory tracks included in the connectivity corridor over the corridor surface (%.km^-2^).

As the proportion of locations included in a corridor depends on the corridor surface, we calculated the ratio of included locations over the surface of the corridor.

The corridor with the largest ratio of locations included in the corridor over the corridor surface was the 99^th^ percentile connectivity corridor. However, the variation of the ratio over the 15 populations was high, suggesting a great heterogeneity in the accuracy of this corridor.

Appendix S12: Results of connectivity modeling. On the left side are represented observed migration routes (spring in green and autumn in brown) and summer and winter ranges (orange and blue) of Alpine ibex. On the right side are displayed the connectivity maps obtained from the “leave 10% of whole data out” dataset. The black lines delineate the connectivity corridors as defined in the ‘representation in corridors’ validation method.

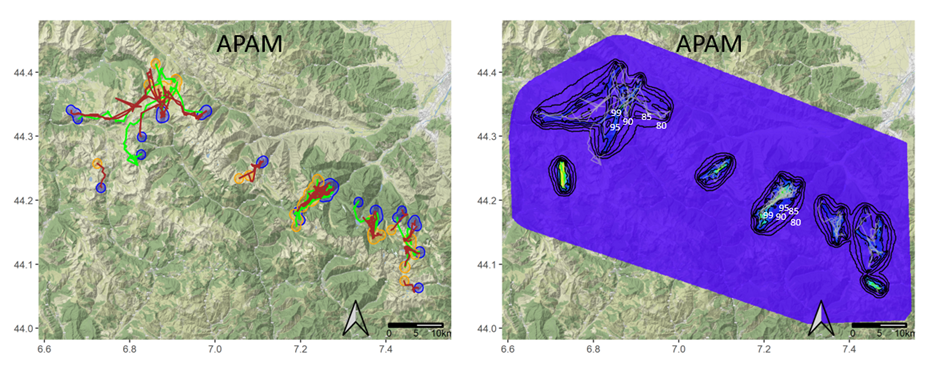

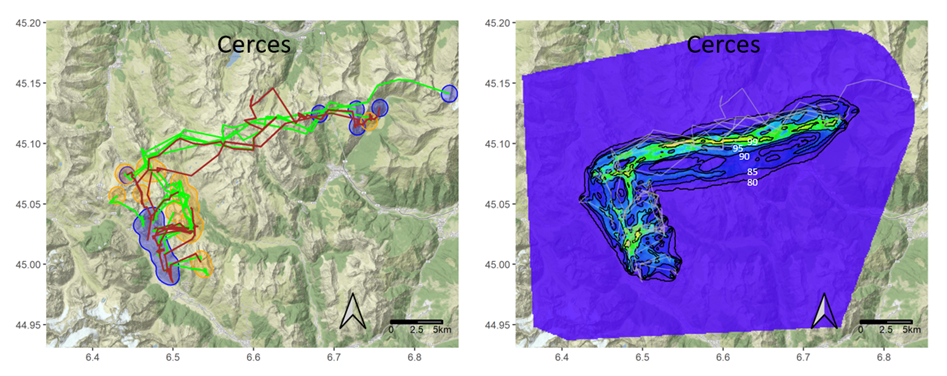

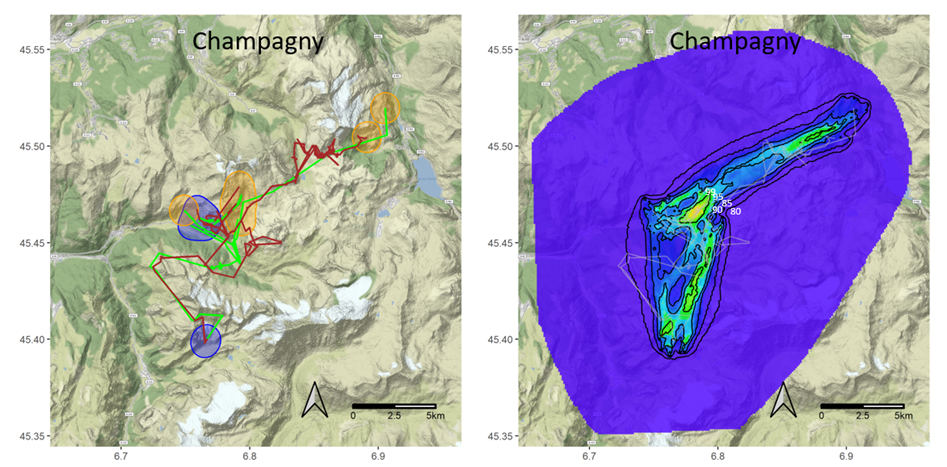

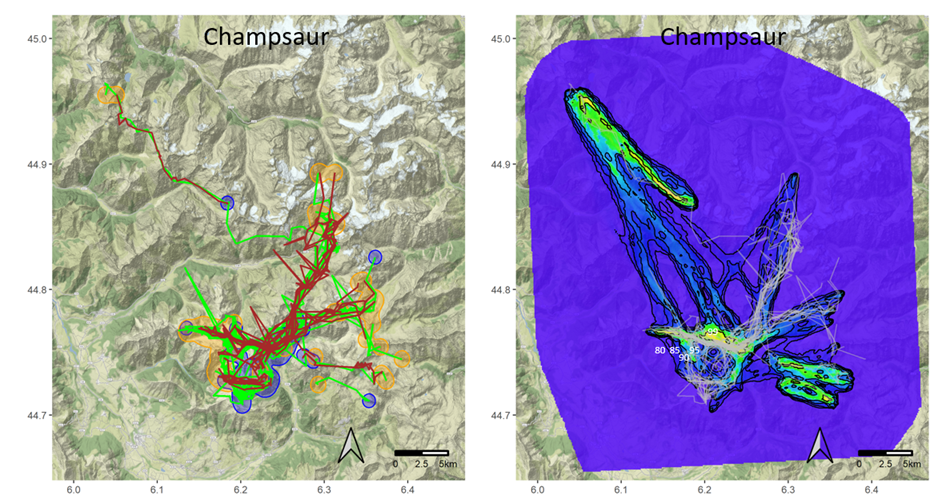

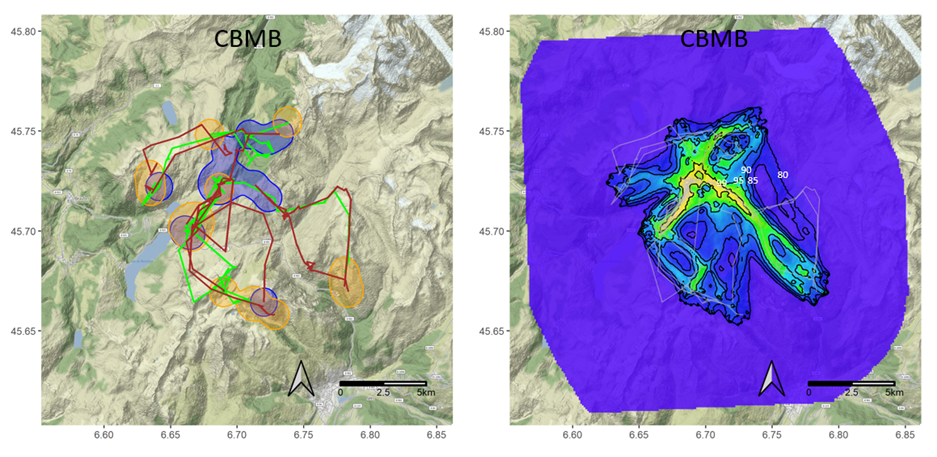

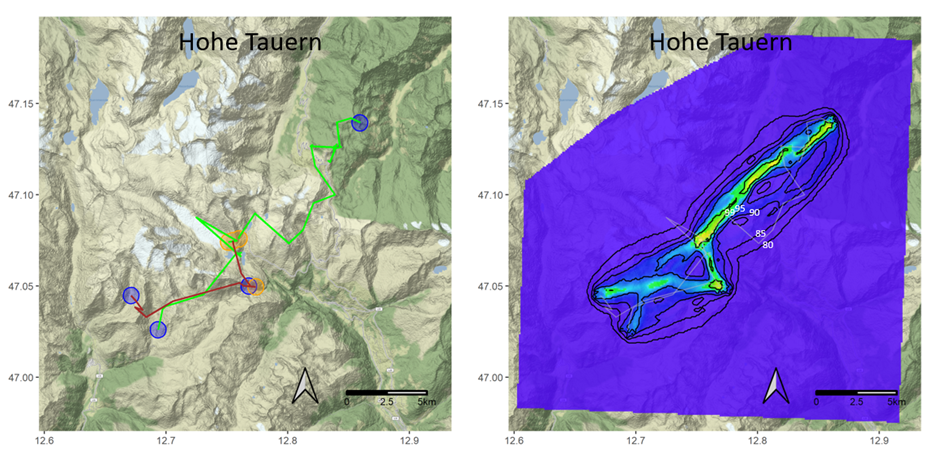

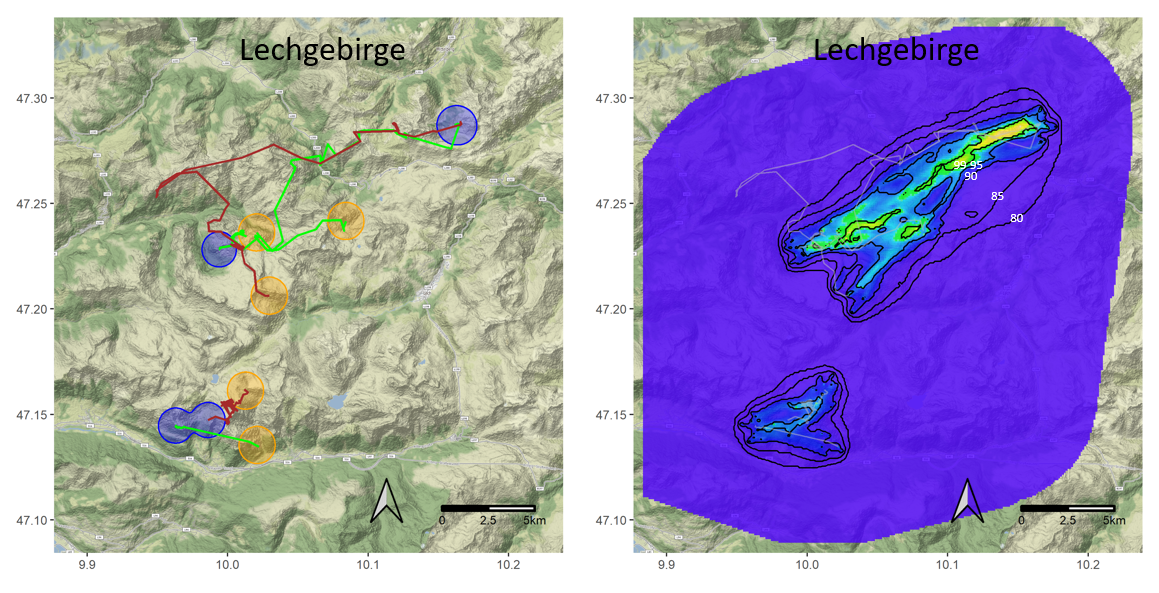

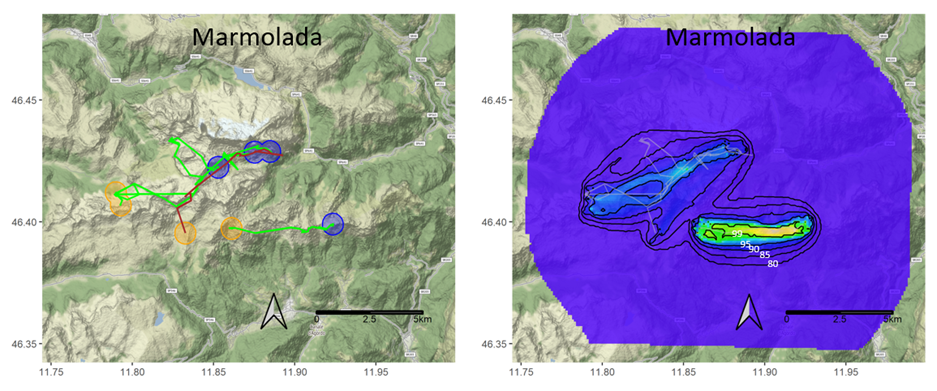

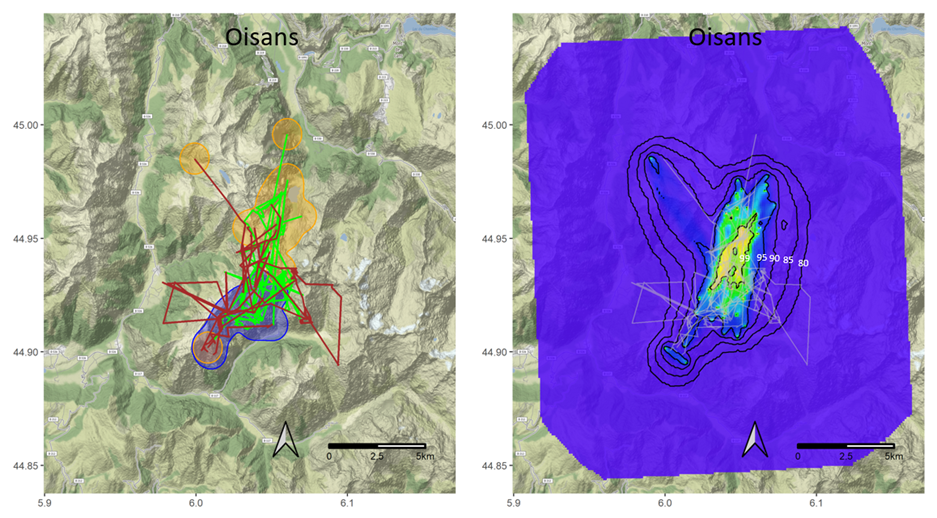

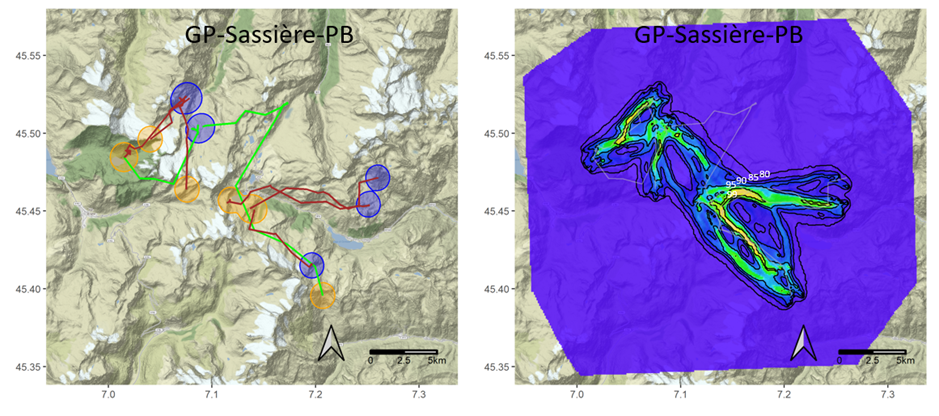

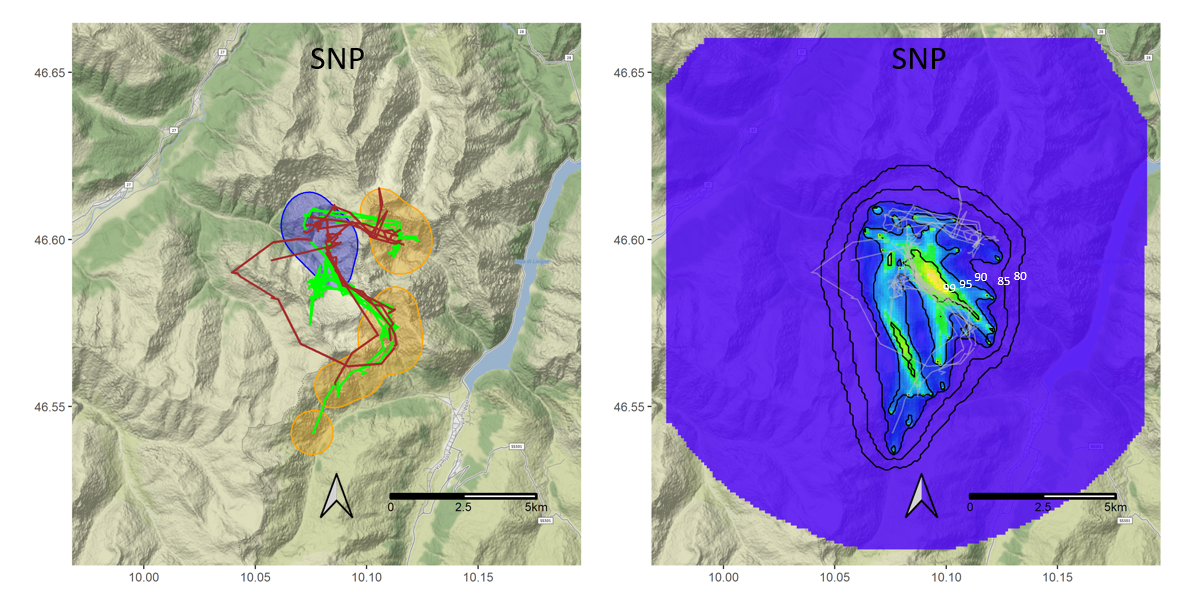

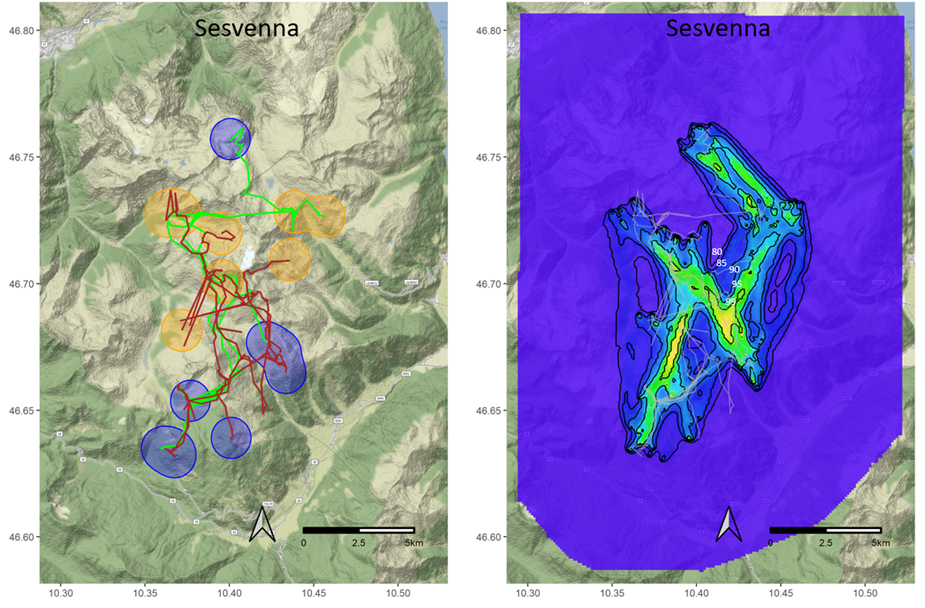

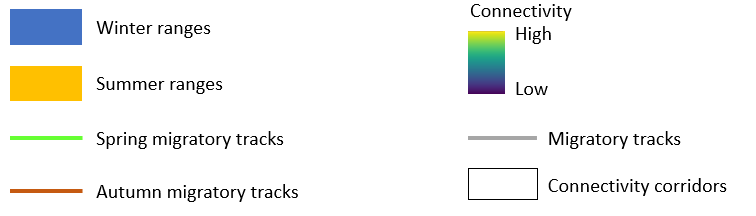

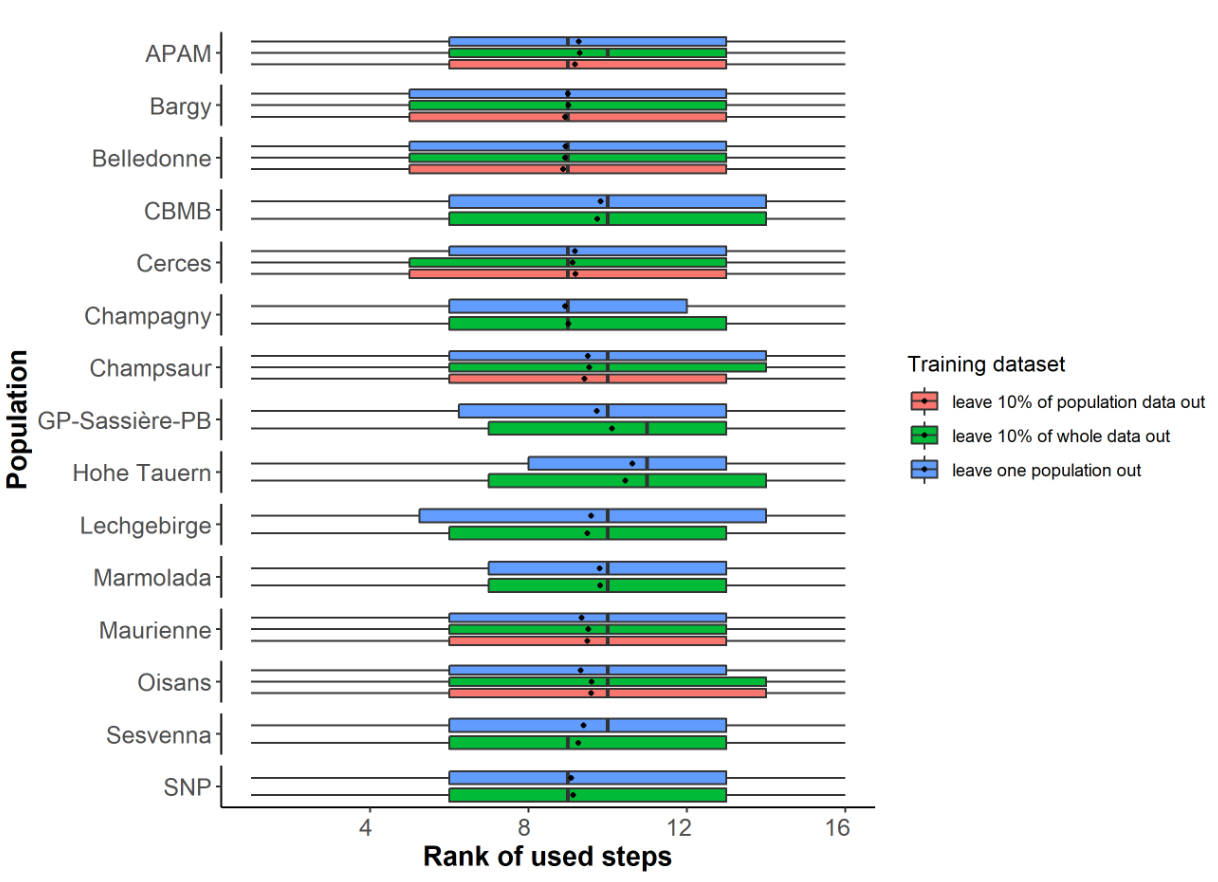
Appendix S13: Results of the ranking validation method. For each population is shown the distribution of the rank of used steps within their strata (one stratum = 1 used step and its matching 15 available steps). Lower and upper limits of the boxes display 1^st^ and 3^rd^ quartiles, the vertical bar and the point respectively display the median and the mean. Leave one population out = connectivity modeled with mean habitat selection of the 14 other populations, leave 10% of whole data out = connectivity modeled with mean habitat selection of all populations, leave 10% of population data out = connectivity modeled with population specific habitat selection preferences.

Appendix S14: Results of the second validation method, representation in corridors (Goicolea et al. 2021).

14-A: Mean proportions (and standard errors) of ibex locations from migratory tracks included in the 80th, 85th, 90th, 95th and 99th connectivity percentile corridors.

| Training dataset | Connectivity corridor (percentile) | | | | |
| --- | --- | --- | --- | --- | --- |
|  | 80 | 85 | 90 | 95 | 99 |
| Leave 10% of whole data out | 93.8(5.0) | 87.8(7.5) | 77.8(11.1) | 53.1(13.7) | 15.7(8.4) |
| Leave one population out | 93.7(6.1) | 88.1(7.8) | 76.8(12.0) | 50.4(14.7) | 15.0(7.6) |
| Leave 10% of population data out | 94.0(5.5) | 88.2(7.5) | 76.9(10.6) | 51.9(11.8) | 14.9(7.1) |

14-B: Proportions of ibex locations from migratory tracks included in the 80th, 85th, 90th, 95th and 99th connectivity percentile corridors for each population and training dataset.

|  |  | Connectivity corridor (Percentile) | | | | |
| --- | --- | --- | --- | --- | --- | --- |
| Population | Training dataset | 80 | 85 | 90 | 95 | 99 |
| APAM | leave 10% of whole data out | 0.969 | 0.962 | 0.883 | 0.718 | 0.241 |
| APAM | leave one population out | 0.971 | 0.962 | 0.878 | 0.705 | 0.230 |
| APAM | leave 10% of population data out | 0.973 | 0.952 | 0.866 | 0.693 | 0.221 |
| Bargy | leave 10% of whole data out | 0.957 | 0.928 | 0.790 | 0.471 | 0.066 |
| Bargy | leave one population out | 0.925 | 0.854 | 0.710 | 0.418 | 0.083 |
| Bargy | leave 10% of population data out | 0.981 | 0.938 | 0.776 | 0.461 | 0.108 |
| Belledonne | leave 10% of whole data out | 0.896 | 0.763 | 0.584 | 0.312 | 0.064 |
| Belledonne | leave one population out | 0.914 | 0.811 | 0.628 | 0.330 | 0.066 |
| Belledonne | leave 10% of population data out | 0.926 | 0.814 | 0.610 | 0.336 | 0.063 |
| Cerces | leave 10% of whole data out | 0.938 | 0.812 | 0.631 | 0.370 | 0.107 |
| Cerces | leave one population out | 0.939 | 0.781 | 0.626 | 0.377 | 0.105 |
| Cerces | leave 10% of population data out | 0.913 | 0.761 | 0.616 | 0.362 | 0.090 |
| Champagny | leave 10% of whole data out | 0.974 | 0.953 | 0.849 | 0.548 | 0.063 |
| Champagny | leave one population out | 0.972 | 0.951 | 0.826 | 0.500 | 0.063 |
| Champagny | leave 10% of population data out | 0.974 | 0.954 | 0.861 | 0.547 | 0.083 |
| Champsaur | leave 10% of whole data out | 0.847 | 0.745 | 0.602 | 0.393 | 0.164 |
| Champsaur | leave one population out | 0.787 | 0.708 | 0.534 | 0.358 | 0.175 |
| Champsaur | leave 10% of population data out | 0.856 | 0.764 | 0.613 | 0.404 | 0.152 |
| CBMB | leave 10% of whole data out | 0.876 | 0.799 | 0.700 | 0.555 | 0.219 |
| CBMB | leave one population out | 0.882 | 0.826 | 0.740 | 0.448 | 0.174 |
| CBMB | leave 10% of population data out | 0.861 | 0.797 | 0.703 | 0.523 | 0.221 |
| Hohe Tauern | leave 10% of whole data out | 0.978 | 0.890 | 0.786 | 0.656 | 0.217 |
| Hohe Tauern | leave one population out | 0.973 | 0.919 | 0.838 | 0.676 | 0.324 |
| Hohe Tauern | leave 10% of population data out | 0.972 | 0.907 | 0.837 | 0.567 | 0.158 |
| Marmolada | leave 10% of whole data out | 1.000 | 0.858 | 0.775 | 0.596 | 0.213 |
| Marmolada | leave one population out | 1.000 | 0.866 | 0.805 | 0.631 | 0.208 |
| Marmolada | leave 10% of population data out | 0.999 | 0.931 | 0.721 | 0.561 | 0.245 |
| Maurienne | leave 10% of whole data out | 0.924 | 0.871 | 0.731 | 0.471 | 0.144 |
| Maurienne | leave one population out | 0.970 | 0.905 | 0.757 | 0.441 | 0.116 |
| Maurienne | leave 10% of population data out | 0.967 | 0.862 | 0.721 | 0.435 | 0.104 |
| Oisans | leave 10% of whole data out | 0.954 | 0.924 | 0.868 | 0.747 | 0.244 |
| Oisans | leave one population out | 0.974 | 0.937 | 0.866 | 0.700 | 0.214 |
| Oisans | leave 10% of population data out | 0.979 | 0.960 | 0.925 | 0.740 | 0.226 |
| GP-Sassière-PB | leave 10% of whole data out | 0.847 | 0.832 | 0.770 | 0.521 | 0.081 |
| GP-Sassière-PB | leave one population out | 0.833 | 0.833 | 0.637 | 0.422 | 0.137 |
| GP-Sassière-PB | leave 10% of population data out | 0.825 | 0.811 | 0.734 | 0.513 | 0.090 |
| Sesvenna | leave 10% of whole data out | 0.968 | 0.921 | 0.834 | 0.573 | 0.162 |
| Sesvenna | leave one population out | 0.977 | 0.962 | 0.851 | 0.521 | 0.199 |
| Sesvenna | leave 10% of population data out | 0.967 | 0.925 | 0.827 | 0.566 | 0.147 |
| SNP | leave 10% of whole data out | 0.990 | 0.988 | 0.956 | 0.694 | 0.323 |
| SNP | leave one population out | 0.989 | 0.987 | 0.958 | 0.728 | 0.076 |
| SNP | leave 10% of population data out | 0.990 | 0.977 | 0.916 | 0.657 | 0.271 |
| Lechgebirge | leave 10% of whole data out | 0.955 | 0.930 | 0.896 | 0.348 | 0.050 |
| Lechgebirge | leave one population out | 0.947 | 0.906 | 0.872 | 0.312 | 0.086 |
| Lechgebirge | leave 10% of population data out | 0.915 | 0.880 | 0.806 | 0.423 | 0.061 |

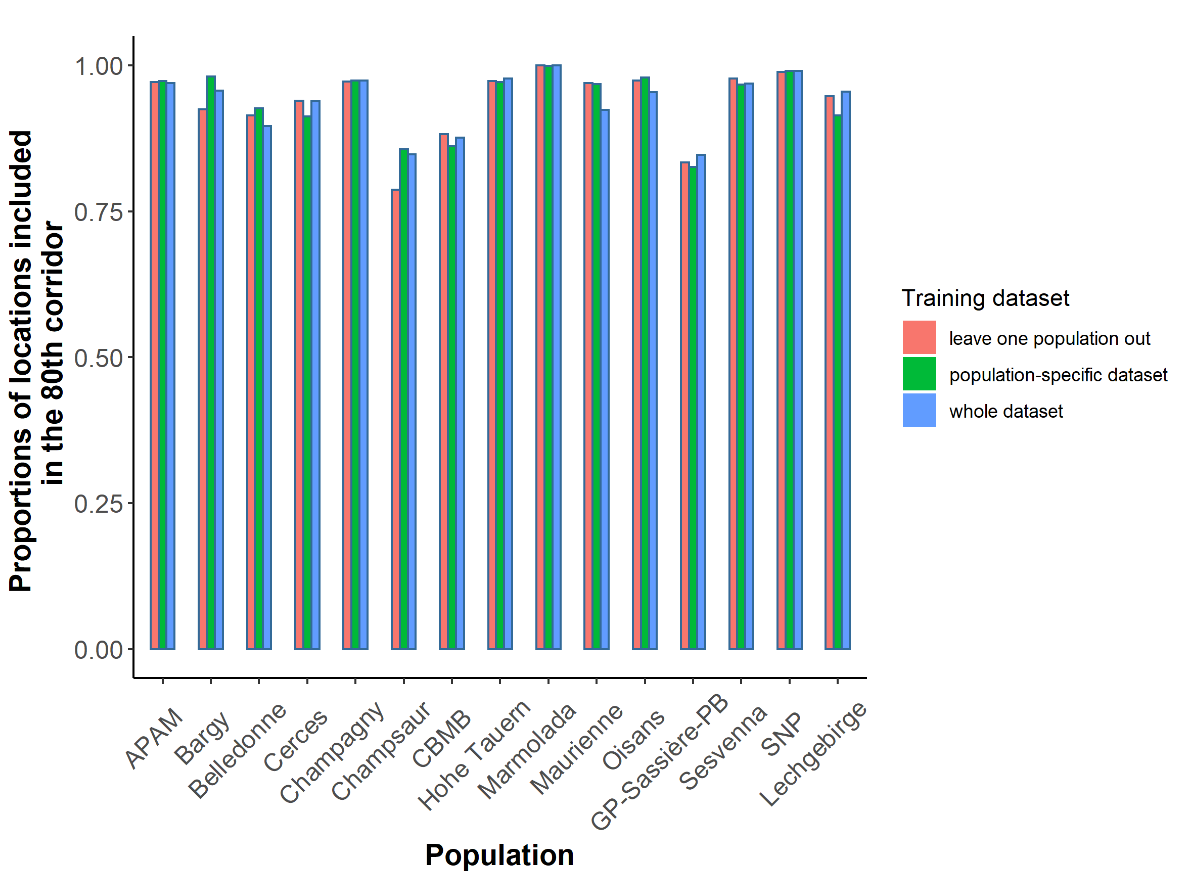
Appendix S15: Representation in corridor validation method. Proportion of ibex locations from migratory tracks included in the 80th, 95th and 99th connectivity percentile depending on the training dataset and the population.

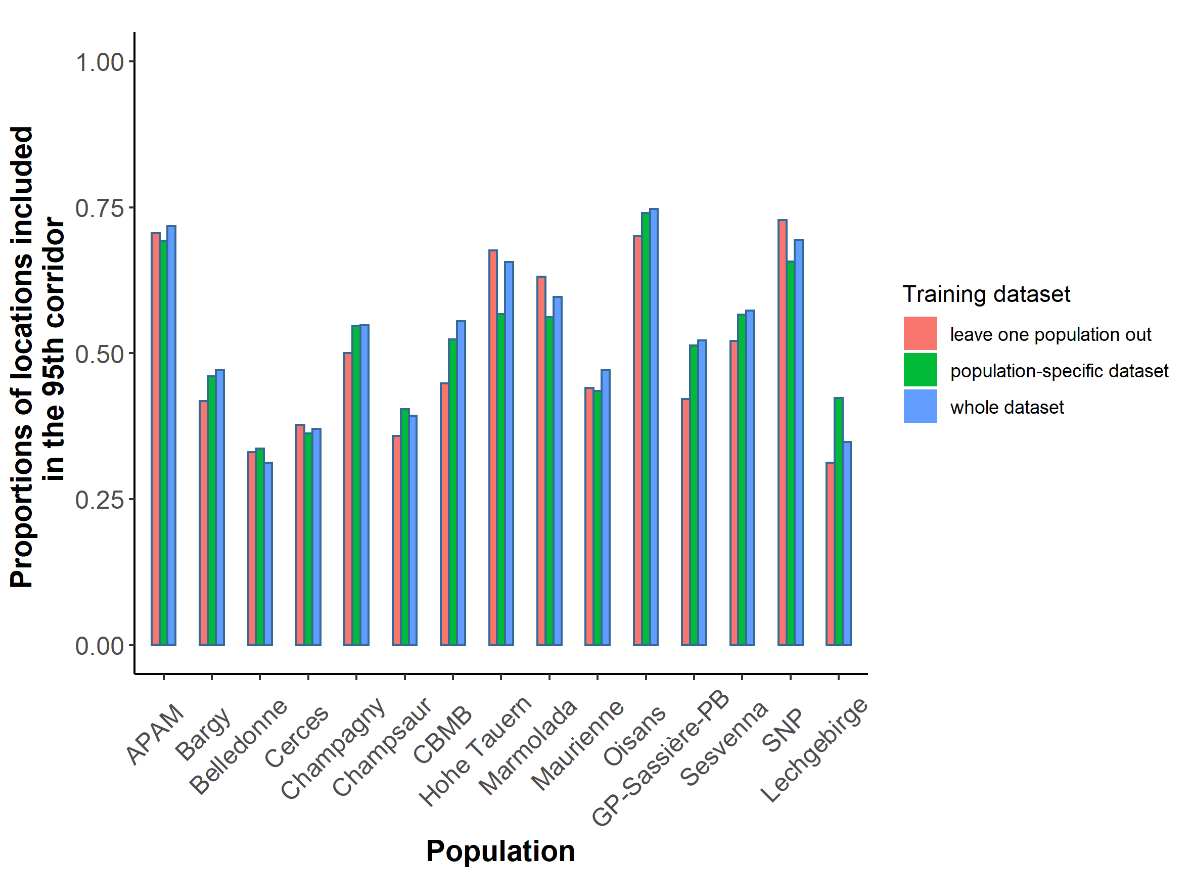

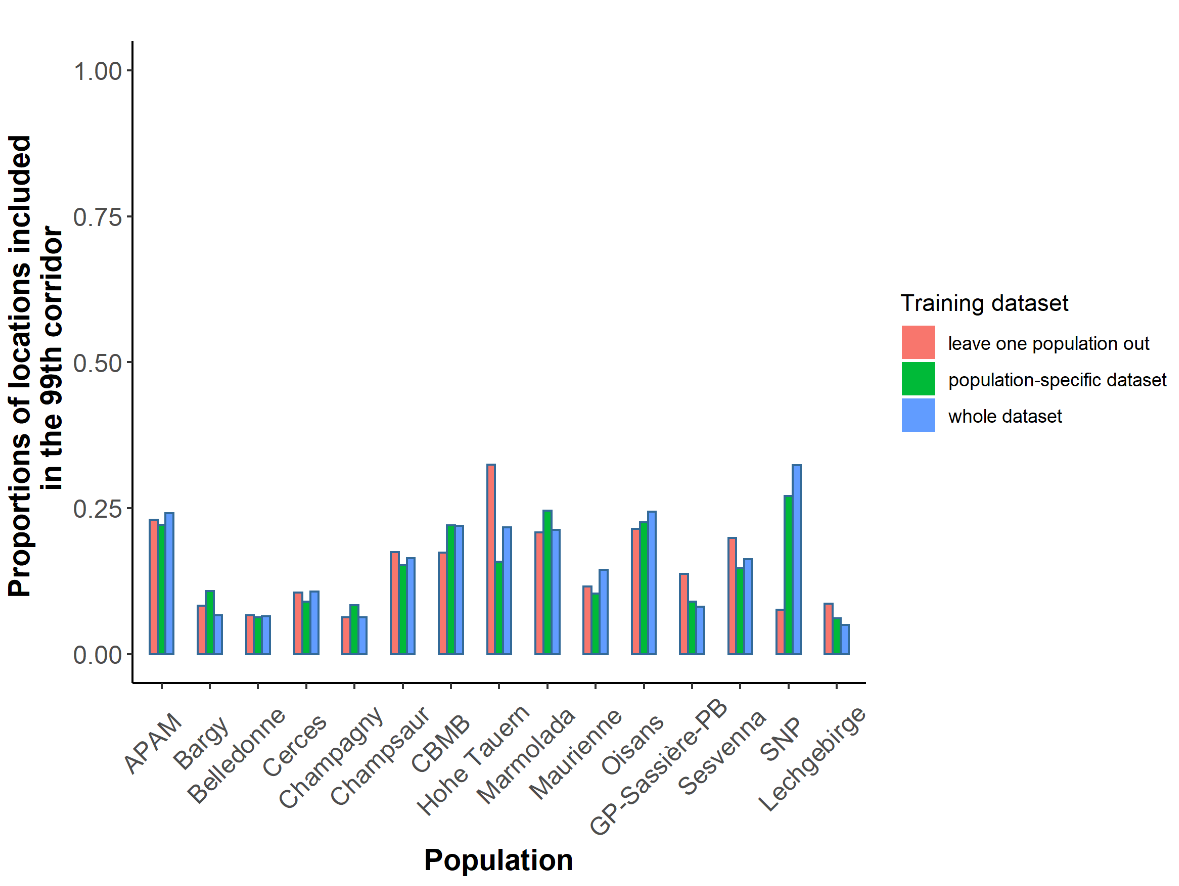
